# BOLD activity during emotion reappraisal positively correlates with dietary self-control success

**DOI:** 10.1101/542712

**Authors:** Silvia U. Maier, Todd A. Hare

## Abstract

We combined established emotion regulation and dietary choice tasks with fMRI to investigate behavioral and neural associations in self-regulation across the two domains in human participants. We found that increased BOLD activity during the successful reappraisal of positive and negative emotional stimuli was associated with dietary self-control success. This cross-task correlation was present in medial and lateral prefrontal cortex as well as the striatum. In contrast, BOLD activity during the food choice task was not associated with self-reported emotion regulation efficacy. These results suggest that neural processes utilized during the reappraisal of emotional stimuli may also facilitate dietary choices that override palatability in favor of healthfulness. In summary, our findings indicate that the neural systems supporting emotion reappraisal can generalize to other behavioral contexts that require reevaluation of rewarding stimuli and outcomes to promote choices that conform with the current goal.

## Introduction

Cognitive strategies and, more recently, the neural mechanisms used to regulate thoughts and actions have been intensely studied in many scientific disciplines. These studies have examined numerous forms of self-regulation, but one prominent strategy is the reappraisal of stimuli encountered in the world (Scherer et al., 2001; Ochsner and Gross, 2005; Etkin et al., 2015). Pioneering studies by Mischel and colleagues (Mischel et al., 1972; Mischel and Moore, 1973; Mischel and Underwood, 1974; Mischel and Baker, 1975) revealed that presenting tempting stimuli as less approachable (e.g., asking participants to imagine food stimuli as abstract pictures) increased the ability to delay gratification (see also Silvers et al. (2014)). Thus, actively reconstructing and reconsidering situations or experiences may enhance control over one’s desires and emotions (Kross et al., 2005; Kross and Mischel, 2010). Converging evidence shows that reappraising stimuli decreases cravings for immediate rewards such as drugs or food when stimuli related to these rewards are viewed (Kober et al., 2010; Hollmann et al., 2012; Hutcherson et al., 2012; Siep et al., 2012; Szasz et al., 2012; Zhao et al., 2012; Giuliani et al., 2013; Yokum and Stice, 2013; Giuliani et al., 2014; Beadman et al., 2015; Svaldi et al., 2015; Boswell et al., 2018; Garland et al., 2018; Reader et al., 2018). Reappraisal appears to be a highly relevant self-regulatory skill. However, although ample evidence shows that individuals can dampen their cravings by reappraising food stimuli, and recent studies show that training to reappraise food stimuli (Boswell et al. (2018)) translates into healthier food choices when participants are asked to decide what to eat at the end of the study, or right after applying reappraisal strategies on each trial (Hutcherson et al. (2012), Schmidt et al. (2018)), it remains unclear whether such regulation skills generalize between task types. For example, does the ability to reappraise emotional stimuli correlate with self-control in eating behavior?

It has been argued previously that self-regulation skills are domain-general, but temptations or challenges may be still be domain-specific (Duckworth and Tsukayama, 2015), potentially requiring very different neural responses. Most of the existing evidence for domain generality or specificity comes from meta-analyses, which either compare different neural candidate mechanisms, or broad classes of tasks using *between-subjects* designs. The degree to which self-regulatory processes in different laboratory tasks and real-life situations share common cognitive and neural substrates is debated (Braver and Barch, 2002; Ridderinkhof et al., 2004; Collette et al., 2006; Dosenbach et al., 2007; Duncan, 2010; Duckworth and Kern, 2011; Heatherton and Wagner, 2011; Tabibnia et al., 2011; Ochsner et al., 2012; Duckworth and Tsukayama, 2015; Kelley et al., 2015; Han et al., 2018; Kragel et al., 2018; Langner et al., 2018; Eisenberg et al., 2019). However, it is important to understand whether an individual can engage neural mechanisms of regulation, be they overlapping or distinct, to modulate behavior to the same relative degree in different behavioral domains. For example, Tusche and Hutcherson (2018) found a significant correlation between the regulation of food and altruistic choices, and Berkman et al. (2011) reported that inferior frontal gyrus (IFG) activity during a go/no go task moderated the relationship between craving and smoking. A recent study by (Suzuki et al., 2020) showed that regulation of alcohol cravings and regulation of negative emotion share a common neural substrate in the left inferior frontal gyrus (ventrolateral prefrontal cortex, vlPFC). Such questions and predictions can be tested directly using *within-subjects* designs. Therefore, here we compare self-regulation in the forms of reappraisal of emotion evoking scenes and health-oriented dietary choices in order to test for associations between these two behaviors.

Meta-analytic evidence suggests that the neural systems supporting the reappraisal of emotions and dietary self-control overlap to some extent (Langner et al., 2018). Previous work looking at emotion regulation has shown that explicit reappraisal recruits prefrontal cortex regions including dorsolateral prefrontal cortex (dlPFC), dorsomedial PFC (dmPFC), ventrolateral PFC (vlPFC), ventral anterior cingulate cortex (vACC), ventromedial PFC (vmPFC), and the supplementary motor area (SMA) (Gross, 1998; Ochsner and Gross, 2005; Wager et al., 2008; Ochsner et al., 2012; Buhle et al., 2014; Etkin et al., 2015; Morawetz et al., 2017a). These regions appear to modulate the reactivity of the insula and dorsal ACC, amygdala and ventral striatum (Delgado et al., 2008; Wager et al., 2008; Etkin et al., 2015; Morawetz et al., 2017b). Similarly, dietary self-control has been reported to involve a set of prefrontal regions including dlPFC, dmPFC, dACC, and vmPFC (Hare et al., 2009; Hare et al., 2011; Harris et al., 2013; Maier et al., 2015; van Meer et al., 2017). However, all of these regions have been reported to be involved in a wide range of behaviors beyond self-regulation, and thus it is unclear what, if any, conclusions we can draw from partially overlapping patterns of activity between emotional reappraisal and dietary choice. Thus, it remains an open question what, if any, neural activation patterns might underlie individual differences in self-regulation success across domains. Therefore, it is important that studies directly test whether neural processes underlying self-regulation within one domain are associated with behavioral outcomes in another domain.

In order to directly compare and contrast neural processing and regulatory success between dietary and emotional self-regulation, we tested the same individuals using both established emotion reappraisal (Ochsner et al., 2002; Wager et al., 2008) and dietary self-control tasks (Hare et al., 2009). For both tasks, we varied the challenge level across trials such that participants faced trials that ranged from small to large challenges. We hypothesized that, if neural activity patterns during the reappraisal of emotional scenes are relevant to or correlated with processes that aid dietary self-control, then individual differences in BOLD activity during successful reappraisal will be associated with success in the dietary self-control task or vice versa.

## Materials and Methods

### Participants

Forty-three healthy adults (18 men) participated in this study. All participants were German native speakers and maintained a health-oriented lifestyle (including a specific interest in healthy eating), but also enjoyed eating snack foods (e.g. chocolate, cake, cookies, chips or crackers) and did so on at least two occasions per week. We used the Beck Depression Inventory I (Beck et al., 1978), German validated version by Hautzinger et al. (1995), and Toronto Alexithymia Scale (Bagby et al., 1994), German validated version by Franz et al. (2008), to screen for depression and emotion blindness because both conditions have been associated with altered emotion perception. All participants provided written informed consent at the day of the experiment according to the Declaration of Helsinki, and the study was conducted in accordance with the regulations of the Ethics Committee of the Canton of Zurich.

Five participants had to be excluded from dietary self-control analyses: two did not complete this task, for one the experiment could not be constructed with a sufficient number of challenging trials, one did not comply with the instructions, and one never chose to eat during the self-control challenge trials. This left a sample of 17 men (mean age = 22.47 ± 2.27 SD years; BMI mean = 22.76 ± 2.34 SD) and 21 women (mean age = 21.5 ± 2.09 SD years; BMI mean = 21.10 ± 2.25 SD) for the behavioral analyses of dietary choices. One additional participant had to be excluded from the fMRI dietary choice analyses for excessive head motion, but this dataset was included in the behavioral analyses. Seven participants were excluded from reappraisal analyses: five fell asleep during a substantial portion of the task (detected by the eye-tracker), one deliberately closed the eyes during negative pictures (reported during debriefing), and one reported experiencing discomfort due to head positioning during the task. We reasoned that the participant who was uncomfortable, but remained in the scanner without complaint until after the study was engaging in constant self-regulation that would interfere with our analyses. One additional woman was excluded from fMRI analyses for this task due to excessive head motion. This left 35 usable fMRI datasets for the reappraisal task and 37 for the dietary self-control task. In total 31 participants (17 women) completed the reappraisal and dietary self-control tasks and had good fMRI data quality during both. We used a priori criteria that are well established in our lab to make our exclusion decisions. All datasets were excluded before analyzing any behavioral data. For recruiting, we followed the previously published cutoffs of our laboratory for this task (Maier et al., 2015; Maier et al., in press): only individuals who consumed snack foods at least on two occasions per week on average for the past four weeks and had an interest in maintaining a healthy lifestyle took part. We only removed participants based on the observation of the experimenter that a given participant clearly did not follow through on the task (e.g., by falling asleep or closing the eyes deliberately), or was experiencing a condition that precluded measuring the emotion task cleanly (one participant only reported after the scan that he was in pain due to the tight fitting headcoil). These observations were recorded in the lab notebook at the time of data collection and these datasets were excluded from the analyses of the respective task a priori. For the fMRI data, we applied quality checks based on the realignment parameters from the preprocessing. In case motion in the X, Y, or Z direction exceeded 2 mm or 2 degrees tilt, we first tried realigning to another trial. We then accounted with a regressor of non-interest for any times for which there was still a deviation greater 2 mm or 2 degrees from the reference slice for short periods (see information in the fMRI analysis section on flagged volumes). For one participant, who for the second half of the food choice run had moved more than 3mm and 2 degrees, we excluded the food choice run due to excessive motion before running any further analyses on the food choice data.

### Procedure

Participants were reminded by email on the day before to their study appointment that on the study day, they should eat a small meal of approximately 400 calories 3 hours before their appointment, and in the 2.5 hours leading up to the appointment should consume nothing but water. Together with the study inclusion, instruction and behavioral ratings that were first completed in the laboratory session, this ensured a fasting period of 3 hours before the dietary choice task. Initially, a 6-minute baseline heartbeat measurement was taken while participants were lying supine in a comfortable position in a quiet room. Participants then rated a set of 180 foods for taste (regardless of healthfulness) and health (regardless of taste) on a generalized visual analog scale with markers in steps of 1 from −5 to +5 (with −5 being not at all, and +5 being maximally healthy / tasty), or vice versa. The middle of the scale showed a zone that was termed “neutral” and comprised the area that corresponded to −5 and +5% of the total scale length centered on zero (Figure 1A). We randomly determined whether participants would use a rating scale in which the left-right orientation ranged from negative to positive, or positive to negative. We ensured that individual participants rated food properties and later feelings using the same directionality.

**Figure 1.**
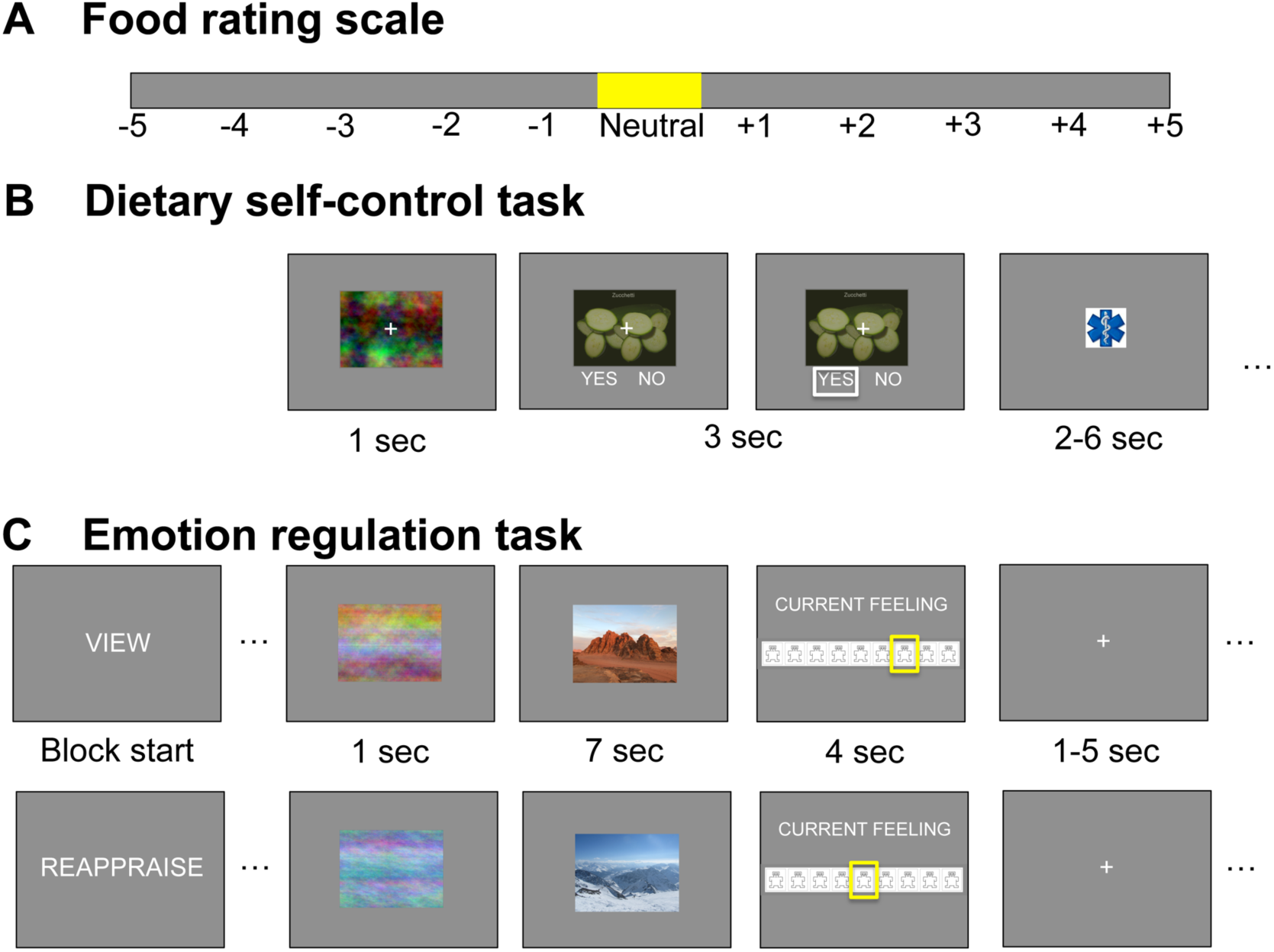
Behavioral tasks: Participants rated 180 food stimuli for taste and health using the rating scale depicted in panel **A**. Items rated “neutral” (falling within ± 5% of the scale around zero) were not presented in the choice set. The order of rating from −5 (“very untasty/unhealthy”) to +5 (“very tasty/healthy”) or vice versa was counterbalanced across participants. In the dietary self-control task (**B**), participants had to choose what to eat at the end of the study. Stimuli were first presented for 1 second as a phase-scrambled image before participants had 3 seconds to choose whether to eat the food by pressing left or right (yes / no, order counterbalanced). The selected option was framed in white for 0.1 seconds. Trials were followed by a jittered 2-6 second inter trial interval. In the emotion regulation task (**C**), participants were presented with positive, negative and neutral stimuli from the International Affective Picture System (IAPS). In blocks of 20 trials, participants were asked to “view” the positive and negative images or to “reappraise” the content such that the elicited feelings got weaker. Neutral images were only presented in the “view” condition. At the beginning of the block, a short verbal instruction for the block appeared for 1 second. An abbreviated reminder (“V” for “view” and “R” for “reappraise”) was then displayed centered on the stimuli instead of the fixation cross. First a phase-scrambled version of the stimulus was presented for 1 second together with the cue. Then the image was revealed for 7 seconds, in which participants had to try and reappraise the content of the picture in order to regulate their feelings or let their feelings evolve naturally. Participants then had 4 seconds to rate their current feeling on a 9-point Self-Assessment-Manikin scale (4^th^ screen). Participants rated both foods and feelings using the same directionality (counterbalanced; from negative to positive or vice versa). Trials were separated by a jittered 1-5 second inter trial interval. After each block of reappraising or viewing, participants were given a 15-second break. Note that in this figure, we have replaced the IAPS stimuli by our own photos for display purposes.

Next, participants received a short practice session to familiarize themselves with the dietary self-control task. At the start, they were reminded to try and choose healthier foods as often as they could, bearing in mind that they would have to eat the item they chose if this trial was drawn to be realized in the end. They made 5 practice choices to get accustomed to the choice screen. The experimenter then introduced the Self-Assessment Manikin (SAM) Scale for rating current emotions according to the procedure detailed in Lang et al. (1999) and explained the reappraisal task using a standardized instruction sheet with one example for positive and negative pictures. Participants were instructed to practice down-regulating their feelings elicited by both negative and positive pictures from the International Affective Picture System by Lang and colleagues. In the *view* condition, they were instructed to watch the presented image and become aware of the feelings that this image evokes. They should not try to alter these feelings. In the *reappraisal* condition, participants should watch the image and try to come up with a different story that could explain the scene, such that the evoked feeling becomes weaker. Negative feelings should become less negative, and positive feelings less positive. For example, one could think of the image as a scene or mock-up from a movie: Things are not as bad or good as they seem, but just staged.

Participants then practiced with a computerized version of the task as it was presented in the fMRI scanner, first for 2 pictures with free timing, and then for 2 pictures with the timing for picture presentation and emotion rating that was applied during the scan. Before going into the scanner, participants rated their current hunger feeling on a visual analog scale with anchors in steps of 1 from −5 to +5 (with −5 being not at all, and +5 being maximally hungry), or vice versa. We transformed these values into percentages of maximal hunger feeling. Across the group, the mean hunger rating was 67% ± 18% SD. In the scanner, participants completed the dietary self-control task and emotion regulation task each in a single run with 100 trials, in counter-balanced order. After the first run, the anatomical scan was collected to allow for a washout period of 7 minutes between the tasks.

After the MRI scans, participants re-rated all 40 stimuli that had been presented in the reappraisal conditions while sitting at a standard computer terminal. Participants were asked to rate the images as in the “viewing” condition, i.e. rating the feeling elicited by the image without altering this emotion.

Lastly, there was a 30-minute waiting period during which one of the food choices was realized for each participant. In case the participant had chosen to eat this food, they were asked to do so within the 30-minute waiting period. If they had refused to eat this food, they were asked to stay in the lab for these 30 minutes without eating anything else. In case the trial was missed, the computer chose randomly whether the participant would have to eat the food or not, thereby incentivizing participants to deliberately make their choices. Participants were fully informed about these procedures before beginning the study. Participants also filled in a battery of psychometric questionnaires during this waiting period. At the end of the 30 minutes, participants were paid a flat fee of 90 CHF for their participation in this 3-hour study.

### Reappraisal task

In the emotion regulation task (Figure 1C), before each block of 20 trials, the condition “view” or “reappraise” was displayed for 1 second. All trials for the respective condition were performed in one block, and participants saw each stimulus only once during the fMRI session. Participants first saw a scrambled version of the stimulus image for 1 second centered on the screen before the stimulus was displayed in the same spot for 7 seconds. During this time, participants had to either passively view the image without altering their feelings, or reappraise their feeling according to the practiced procedure so that their feelings became weaker. To remind them of the condition to be applied, a shortened cue (“V” for view or “R” for “reappraise”) replaced the fixation cross on top of the stimulus. We omitted the letters in the figure for clarity. Participants then had 4 seconds to rate their current feeling on a 9-point SAM valence scale. A jittered inter-trial interval (uniformly sampled from 1 to 5 seconds) separated one trial from the next.

Block types (Reappraise Positive, Reappraise Negative, View Positive, View Negative, View Neutral) were presented in 5 different orders that were pseudo-randomized across participants. Each block was followed by a 15-second break (with the word “pause” appearing over a countdown that showed the remaining seconds of break time).

IAPS Stimuli were selected based on a validation study in a German-speaking sample of young adults (Grühn and Scheibe, 2008). Based on the mean ratings given by young adults in this dataset, we identified 40 images that scored highest on positive and 40 images that scored highest on negative valence, skipping any that showed foods, and proceeding to the next best-scoring images as a replacement. We distributed the positive and negative images each into two sets such that both sets in each domain were equated on average for arousal (mean negative: 6.99 ± 0.44; mean positive: 2.86 ± 0.43 based on the ratings of the sample in Grühn and Scheibe (2008)). We randomly allocated for each of our participants which set they would see in the “view” and “reappraise” condition. We then identified 20 images that scored neutral on both valence and arousal.

### Dietary self-control task

In the dietary self-control task (Figure 1B), participants were shown one food in the center of the screen on each trial, and had to indicate within the 3-second response window whether they wanted to eat this food or nothing at the end of the study. Choices were customized based on the input of the participant who, before the food choice task, gave their individual taste and health ratings for a large set of foods. The customized food sets were created such that each participant would face approximately 75 percent challenging choices, in which the presented food was either subjectively i) palatable and unhealthy (i.e. rated above and below the neutral zone on the taste and health rating scales, respectively; see Figure 1A, yellow zone), or ii) healthy and unpalatable. In the remaining choices, health and taste were aligned, so the food was rated as palatable and healthy, or unpalatable and unhealthy. Trial types were randomly intermixed, and a jittered inter-trial interval (uniform draw of 2 to 6 seconds) separated each trial.

### Psychometric inventories

The psychometric questionnaire battery included the Three Factor Eating Questionnaire (Stunkard and Messick (1985), German validated version by Pudel and Westenhoefer (1989)), Dutch Eating Behavior Questionnaire (Van Strien et al. (1986), German validated version by Nagl et al. (2016)), PANAS (to describe their mood for the last week; Watson et al. (1988), German validated version by Krohne et al. (1996)), BIS-BAS (Carver and White (1994), German validated version by (Strobel et al., 2001)), BIS-15 ( Meule et al. (2011), German validated version of the Barratt Impulsiveness Scale (Patton et al., 1995) short form BIS-15 by Spinella (2007)), and NEO-FFI (Costa and McCrae (1989), German validated version by Borkenau and Ostendorf (2008)).

### fMRI data acquisition

The MRI data were recorded using a Philips Achieva 3 T whole-body scanner with an eight-channel sensitivity encoding head coil (Philips Medical Systems). Stimuli were presented with the Psychophysical Toolbox Software (Psychtoolbox 3.0, Brainard (1997), RRID:SCR_002881) via back-projection to a mirror mounted on the head coil.

We acquired gradient echo T2*-weighted echo-planar images (EPIs) with blood-oxygen-level-dependent (BOLD) contrast (37 slices per volume, Field of View 200 x 132.6 x 200 mm, slice thickness 3 mm, 0.6 mm gap, in-plane resolution 2.5*2.5 mm, matrix 80*79, repetition time 2344 ms, echo time 30 ms, flip angle 77°) and a SENSE reduction (i.e. acceleration) factor of 1.5. Volumes were acquired in axial orientation. We collected 354 volumes during the dietary choice run (∼ 12 minutes), and 679 volumes during the emotion regulation run (∼ 25 minutes). There was one run of data collection for each task. Both runs were collected in ascending order. Before each run, five “dummy” volumes were collected to allow for stabilization of the magnetic field. A T1-weighted turbo field echo structural image was acquired in sagittal orientation for each participant between the functional scans (181 slices, Field of View 256 x 256 x 181 mm, slice thickness 1 mm, no gap, in-plane resolution 1*1 mm, matrix 256*256, repetition time 8.3 ms, echo time 3.89 ms, flip angle 8°). To measure the homogeneity of the magnetic field we collected B0/B1 maps before the first run and before acquiring the structural scan (short echo time = 4.29 ms, long echo time = 7.4 ms). We measured breathing frequency and took an electrocardiogram with the in-built system of the scanner in order to correct for physiological noise.

### fMRI preprocessing

Functional data were spatially realigned and unwarped with statistical parametric mapping software (SPM12, Update Rev. Nr. 6906; Functional Imaging Laboratory, University College London, RRID:SCR_007037), slice-timing corrected, coregistered to the participant’s T1-weighted high resolution structural image and normalized to the individual mean EPI template before segmenting according to the individual T1 scan and smoothing with an isometric Gaussian kernel (4 mm full width at half maximum). In order to account for fluctuations in the BOLD signal due to physiological noise, we finally used RETROICOR as implemented in the TAPAS PhysIO toolbox (Version 2015; open source code available as part of the TAPAS software collection: http://translationalneuromodeling.org/tapas/) by Kasper et al. (2017) to model respiration and heartbeat (Glover et al., 2000; Hutton et al., 2011). Following Harvey et al. (2008), the algorithm implemented in the PhysIO toolbox uses Fourier expansions of different order to the estimate the phases of cardiac pulsation (3rd order), respiration (4th order) and cardio-respiratory interactions (1st order).

### Experimental Design and Statistical Analysis

We sought to identify whether neural processes occurring during reappraisal were associated with the behavioral outcome of another, distinct self-regulation task: dietary self-control. All correlations reported in this paper were calculated using a Bayesian estimation procedure (Kruschke, 2015), where we calculated the Bayesian equivalent of Pearson’s (linear) or Spearman’s (rank) correlation coefficients across all participants.

Our hypothesis was that neural activity during reappraisal would be correlated with dietary self-control success, and potentially vice versa. Note, however, that these two relationships are distinct and a relationship in one case does not indicate or require the other. To compare both reappraisal and dietary self-control abilities, we chose a within-subject design. Based on prior reports of these self-regulation tasks in the literature, we expected a moderate effect size (Webb et al., 2012).

All behavioral analyses presented in this paper were performed with the R (“R Core Team,” 2015), version 3.5.1, RRID:SCR_001905, STAN (Carpenter et al., 2016) and JAGS (Plummer, 2003) statistical software packages. For all Bayesian modeling analyses, we used the default, uninformative priors specified by the brms (Bürkner, 2017) and BEST (Kruschke, 2013, 2015) R-packages, which means that our Bayesian analyses would give very similar results to their frequentist analogs.

SPM12 (Penny et al. (2006), update 6906) was used to preprocess fMRI data and calculate first-level models. FSL’s Randomise tool (Winkler et al., 2014) was used to run nonparametric permutation tests (n = 5000 permutations) with threshold-free cluster enhancement (TFCE) on the group level. We chose to switch to the implementation in FSL 5 (RRID:SCR_002823) for this analysis, because the TFCE and permutation algorithms were more fully documented and computed faster in FSL compared to SPM12.

Figures 4 to 7 were created using the MRIcron and MRIcroGL software packages (http://www.mccauslandcenter.sc.edu/mricro/mricron/, http://www.mccauslandcenter.sc.edu/mricro/mricrogl/, RRID:SCR_002403). Anatomical labels for the tables were derived from the Harvard-Oxford cortical and subcortical atlases (Desikan et al., 2006, RRID:SCR_001476) with FSL’s atlasquery and cluster commands.

In the main text we report T and p values for the strongest contiguous cluster in each analysis. Exact T values at the voxel-level can be found in a Neurovault repository (link: https://www.neurovault.org/collections/YPGQPMUT/). For non-significant contrasts we report the minimum whole-brain corrected p-values (or minimum small-volume corrected p-values where indicated). All analysis code and raw data for the behavioral results can be found at https://github.com/silvia-maier/Maier_Hare_Emotion_and_dietary_selfregulation. Raw fMRI data will be accessible after publication on https://openneuro.org.

The mask for small-volume corrections in the left prefrontal cortex was built from the Harvard-Oxford Cortical Atlas and comprised the frontal pole, inferior frontal gyrus pars operculum and pars triangularis, as well as medial and superior frontal gyrus (14215 voxels).

### Behavioral Analyses

*Reappraisal task.* In the emotion paradigm, reappraisal success was measured as the difference between emotion ratings given when reappraising the image inside the scanner and post-scan ratings made when viewing the same picture again without reappraising it as in Ochsner et al. (2002). We calculated success scores for negative-valence stimuli as the difference, Reappraisal minus View, because the reappraised rating should be higher (i.e. more positive) than the unregulated viewing rating if reappraisal of negative stimuli was successful. The difference, View minus Reappraise, was calculated for positive reappraisal trials, because for positive stimuli the unregulated View ratings should be higher than the reappraised rating when successfully modulating positive emotions. Our primary measure of reappraisal success, the overall emotion reappraisal success score, was computed across both negative and positive images as the mean of the positive plus negative reappraisal success scores. However, we also computed and checked the reappraisal success scores for each valence separately in some cases noted below. We also checked that positive and negative reappraisal success did not differ significantly (see Supplementary Methods, Results, Table S1 and Figure S1).

**Table 1.** Emotion ratings by condition.

| **A. Emotion ratings** |  |  |  |
| --- | --- | --- | --- |
| **Condition** | **Mean rating** | **Standard Deviation** |  |
| Negative View | 2.69 | 0.54 |  |
| Negative Regulate | 4.25 | 0.81 |  |
| Neutral View | 5.26 | 0.41 |  |
| Positive Regulate | 5.21 | 0.90 |  |
| Positive View | 7.09 | 0.67 |  |
| **B. Regression results** |  |  |  |
| **Fixed effects** | **Beta estimate** | **Standard Deviation** | **95% Highest Density Interval** |
| (Intercept) | 5.26 | 0.07 | [5.13; 5.40] |
| Negative View | -2.57 | 0.10 | [-2.77; -2.38] |
| Negative Regulate | -1.00 | 0.15 | [-1.31; -0.70] |
| Positive Regulate | -0.06 | 0.17 | [-0.39; 0.26] |
| Positive View | 1.82 | 0.13 | [1.57; 2.07] |
| Bayesian $R^2$ | 0.65 | 0.01 | [0.64; 0.66] |

**A)** Mean and standard deviation for the emotion ratings given in each block type.
**B)** Emotion ratings were modeled in a Bayesian linear regression model (specified in Eq. 1) as a function of block type, allowing participant-specific random intercepts and participant-specific random slopes. Block type was a factor with five levels (negative view, negative regulate, neutral view, positive regulate, positive view). The results show differences in the ratings with regard to neutral viewing as the baseline condition. The analyses in both A) and B) comprised N = 36 participants.

To test whether ratings differed significantly between the conditions, we conducted the following linear regression:

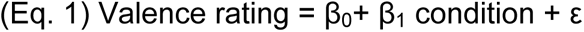

In this model, *valence rating* was the rating given on the respective trial, coded from 1 (very sad) to 9 (very happy) in steps of 1, according to the Self-Assessment Manikin scale, and *condition* was a factor with 5 levels (0 = neutral view, 1 = negative view, 2 = negative reappraisal, 3 = positive reappraisal, 4 = positive view). The model included subject-specific random intercepts and slopes for the condition.

#### Dietary self-control task

In the dietary self-control paradigm, challenging trials were defined as those trials in which health and taste attributes were not aligned. The overall self-control success level was measured as the proportion of all challenging trials in which participants refused to eat a tasty, unhealthy food, or accepted eating a healthy, unpalatable food as in Hare et al. (2009). We tailored each participant’s food choice set such that s/he would face approximately 75 self-control challenges (in which health and taste were not aligned) out of 100 decisions. To classify these challenges, we used the individual health and taste ratings that each participant had given previously on this day for the foods. The number and types of challenges we could present each individual depended on their ratings for the full set of 180 foods. Most participants faced more self-control challenges for items that were unhealthy and tasty (out of 100 choices: minimum 14, median 52.5, maximum 77) than challenges including healthy but unpalatable items (minimum 0, median 15, maximum 46).

To characterize dietary choice patterns, we modeled participant’s choices of the healthier item as a function of taste and health properties with a Bayesian mixed logistic regression model (Eq. 2):

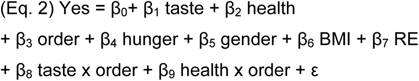

In this model, *Yes* was a binary indicator for choices taking the value 1 when the participant chose to eat the presented item and 0 otherwise, and *taste* and *health* denoted the respective ratings for the item depicted on the screen that were standardized and mean-centered across all participants. The model included subject-specific random intercepts and subject-specific random slopes for the *taste* and *health* attributes, allowing both variables to have differential effects in each participant. To check the robustness of our results, we also included control variables for the main effect of the *order* in which reappraisal and dietary self-control tasks were performed and the interactions of task order and taste and health attributes, as well as the main effects of *hunger* level (in percent, indicated on a visual analog scale from 0, not at all, to 100, maximally hungry), *gender* (male / female, self-reported), Body Mass Index (*BMI*) and restrained eating score (*RE*) on the restraint subscale of the Three Factor Eating Questionnaire (Pudel and Westenhoefer, 1989). Task order and gender were modeled as factors, and standardized scores were used for eating restraint, BMI and hunger level. We chose to include the restrained eating subscale of the TFEQ based on our prior work (Maier and Hare, 2017), in which we showed that RE explained individual variation in dietary self-control behavior beyond the effects of task features.

To test for the determinants of self-control in challenging trials, in which health and taste aspects were not aligned, we modeled self-control success as a function of taste, health and challenge type:

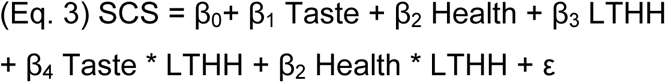

Where *SCS* was a binary variable taking the value of 1 if participants succeeded on this trial and 0 if they did not, *Taste* and *Health* described the within-participant z-scored taste and health ratings for the depicted food, and *LTHH* was a factor with two levels (coded as 1 if participants saw a low-taste/high-health food on this trial and 0 otherwise, i.e. using high-taste/low health challenges as reference). The model included subject-specific random intercepts and subject-specific random slopes for taste, health, challenge type and their interactions.

To test for reaction time differences as a function of trial type, we fit the model described in Eq. 4:

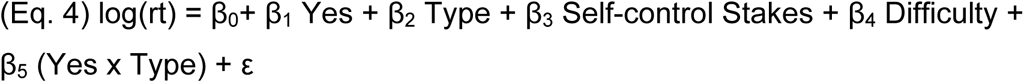

where *log(rt)* was the log-transformed reaction time (rt) for food choices on each trial, *Yes* was a binary indicator for the choice made, equaling 1 if the participant chose to eat the item on the screen and 0 otherwise, and *Type* was a factor with three levels indicating the type of trial (0 = no challenge trials, 1 = challenge trials with high-taste/low-health (HTLH) foods, and 2 = challenge trials with low-taste/high-health (LTHH) foods). The variable *Stakes* was calculated for each trial as described in Eq. (5) and the variable *Difficulty* was calculated for each trial as described in Eq. (6) below. The model included subject-specific random intercepts and subject-specific random slopes for answer, trial type and their interaction and for the self-control stakes and difficulty.

Inspecting the results of fMRI model GLM-SCS (described below) prompted us to investigate more carefully how individuals solved self-control challenges. To this end, we performed an exploratory analysis to investigate how participants tracked the objective challenge and importance of self-control choices. We constructed a measure we call the self-control *stakes* (see Figure 7A, upper left and lower right quadrant). The stakes variable is a combination of the absolute magnitudes of two food attributes: One is the taste of the food, which determines how much taste temptation participants have to resist, or how much aversion they have to overcome in order to eat an unpalatable item. The other, separate aspect is the health benefit or cost they accrue in doing so. The stakes are high both when a very tasty temptation carries with it large health drawbacks (upper left quadrant) and when a highly unpalatable food would yield high health benefits (lower right quadrant). By definition, self-control is only required when the taste and healthiness attributes are in opposition and, therefore, the stakes are zero throughout both the lower left and upper right quadrants. In our analyses, we compute what is at stake in each self-control challenge trial by adding up the absolute value (i.e., the distance from zero, which is in our case equals neutral on the rating scale) of the taste (tr) and health (hr) aspects for all foods in the upper left or lower right quadrants of Figure 7A, according to Eq. 5:

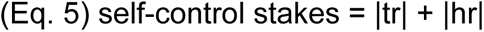

Note that this measure is different from subjective difficulty or decision conflict, which increases the closer weighted taste and health values are to zero (Figure 7B). We calculated the subjective difficulty or decision conflict on each trial according to equation 6:

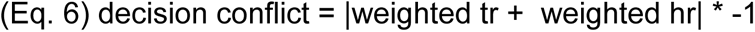

We also sought to estimate the weights on taste and health ratings that capture the subjective importance of taste and health aspects to the decision maker for use in our fMRI analyses. We estimated these weights using the logistic regression model described in Eq. 7 that was calculated for each participant:

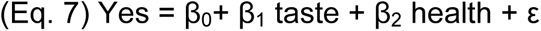

Similar to the model in Eq. 2, *Yes* was a binary indicator for choices taking the value 1 when the participant chose to eat the presented item and 0 otherwise, and *taste* and *health* denoted the respective ratings for the item depicted on the screen that were mean-centered before entering the regression.

### fMRI analyses

#### General linear models

All fMRI models included nuisance regressors for head-motion and cardiac and respiratory effects on each trial. Additionally, in case motion exceeded 2 mm or 2 degrees tilt, a binary regressor flagged this trial, the three preceding trials and one subsequent trial to account for any variance associated with the excessive motion. In total, 12 out of 35 included emotion reappraisal datasets contained flagged volumes (mean = 3.7%, range = [0.7%; 14.1%] of all acquired volumes), and 2 out of 37 included dietary choice datasets (mean = 7.2%, range = [1.4%; 13%]).

In the model of the reappraisal task, onsets for the cue and reappraisal/view screens were modeled as boxcar functions with a duration equaling the cue depiction and task periods (1 and 7 seconds), and rating periods were modeled as boxcar functions with durations equaling the reaction time for the rating. In the fMRI models of the dietary self-control task, regressors were defined as boxcar functions with durations equaling the reaction time on each trial.

#### Correction for multiple testing

We conducted two analyses examining associations between BOLD activity and performance across tasks. Therefore, we applied a Bonferroni-correction to the results resulting in a significance threshold of p < 0.025 for our whole brain analysis. We also conducted a region of interest (ROI) analysis in five regions that have previously been found to be involved in reappraisal processes as well as decision-making (amygdala, dlPFC, hippocampus, striatum, vmPFC) to test whether activity change there in reappraisal success compared to viewing stimuli related to self-control success in the dietary domain. We used a Bonferroni-correction to account for testing in 5 separate regions (resulting significance threshold = p < 0.01).

#### Reappraisal task

Our main general linear model on emotion regulation (GLM-ER) tested for BOLD activity correlated with stimulus reappraisal. GLM-ER modeled events of interest for (1) positive view, (2) positive reappraisal success, (3) positive reappraisal failure, (4) negative view, (5) negative reappraisal success, (6) negative reappraisal failure, (7) neutral view trials, as well as (8) the time during which participants gave their emotion ratings. None of these had any parametric modulators. We calculated a first-level contrast for reappraisal success subtracting BOLD activity during viewing, collapsed over positive and negative modalities. On the group level, we then examined with this contrast whether we detected increases in BOLD activity during Reappraisal Success compared to Viewing, and whether these differential increases for each participant correlated with their empirically measured dietary self-control success level. In addition, to test whether BOLD activity differed for the success in negative compared to positive regulation trials, we calculated the contrasts Positive Reappraisal Success > Negative Reappraisal Success and Negative Reappraisal Success > Positive Reappraisal Success on the individual and group level. For comparison purposes and facilitating further meta-analyses, we also calculated the contrasts Positive Reappraisal > Positive View, Negative Reappraisal > Negative View, and Reappraisal > View that can be inspected in the Neurovault collection, but are not further interpreted here.

#### Dietary self-control task

To assess neural activity during dietary choice, we first calculated GLM-FC (food choice). It modeled events of interest for all trials in which a choice was made. The model included parametric modulators for the subjective food value (linear and quadratic effects), which were orthogonalized. We calculated participant-level and group-level contrasts for the parametric effects of subjective food value.

Subjective food value was calculated as in Maier et al. (2015) and Maier and Hare (2017): We first estimated the logistic regression model specified in Eq. 7 for each participant to model their food choices as a function of taste and health ratings. We then used these taste and health weights that characterize the subjective importance the participant placed on taste and health aspects to weight taste and health ratings for the food choices on each trial and summed up the weighted taste and health values into an overall subjective food value on each trial.

In order to track taste and health aspects separately, we next calculated GLM-TH (taste/health). It modeled events of interest for all trials in which a choice was made. The model included parametric modulators for the taste and health ratings (linear effects), which were not orthogonalized. We calculated participant-level and group-level contrasts for the parametric effects of taste and health.

We calculated a further GLM to test if any brain regions showed differential activity during self-control success versus failure (GLM-SCS), and an exploratory GLM to test whether the brain tracked the stakes of engaging self-control from trial to trial (GLM-ST).

GLM-SCS was constructed after Hare et al. (2009). It modeled events of interest for (1) Self-control success, (2) Self-control failure, (3) Trials without a self-control challenge, and (4) Missed trials. None of these regressors had parametric modulators. Following the analysis of Hare and colleagues, we calculated a second-level correlation of the individual overall self-control success levels with the Self-control Success > No Challenge contrast to track individual differences in the BOLD signal relating to differences in self-control usage. To test for a link with individual differences in emotion reappraisal success, we additionally calculated a correlation of the Self-control success > No challenge contrast with the overall emotion reappraisal success score. Lastly, we also calculated the contrast for Self-control Success > Failure on the individual and group levels.

The exploratory model GLM-ST tests for brain areas correlating with our novel measure of self-control *stakes*, which should be represented in self-control challenges regardless whether or not participants succeeded in using self-control (see Figure 7A, upper left and lower right quadrant). To examine our neural data, we used the stakes measure in addition to decision conflict as parametric modulators in GLM-ST. This GLM included onsets for (1) all trials in which participants decided on palatable-unhealthy or unpalatable-healthy items, (2) palatable-healthy or unpalatable-unhealthy foods, and (3) missed trials. The “stakes” modulator was orthogonalized with respect to decision conflict in order to obtain a readout of the unique signal associated with the need for self-control beyond that correlated with decision conflict (Mumford et al., 2015). The modulator thus explained unique variance associated with the self-control need, adjusted for the variance explained by decision conflict and the variance shared between both. We calculated participant- and group-level contrasts for the stakes parametric modulator in the challenging trials. Note that this model contained also second-order expansions for the parametric modulators in order to control for non-linear effects, but these did not explain any variance, indicating only linear effects were present.

To validate our novel stakes measure, we also re-analyzed a previously acquired dataset with GLM-ST. For the description of this dataset please see our prior reports in Maier et al. (2015) and Maier and Hare (2017). GLM-ST was run including all 51 participants. Note that in this replication test, we only included first-order polynomial expansions in the model given that there were no second-order effects in the original test on the current food choice data.

#### Links between the neural activities in both tasks

We chose to examine the link between the tasks through the Reappraisal Success > View and Self-Control Success > No Challenge contrasts, because these provide a clear interpretation that mechanisms associated with both contrasts also contributed to self-regulation success in both cases.

## Results

### Behavior

#### Behavioral results within each separate task

We found that participants were able to both regulate their emotions and use dietary self-control to select healthier foods well within each experimental task, respectively.

##### Reappraisal task

In the emotion regulation task, we asked participants to either 1) simply view and react naturally, or 2) reappraise photographs with different emotional valence. After seeing or reappraising the pictures for seven seconds, they rated their current affective state using the SAM scale on which 1 indicated the most negative and 9 the most positive emotional valence (Figure 2). To test whether our paradigm was effective, we estimated a Bayesian linear regression that modeled emotion ratings as a function of block type (see Eq. 1 and Table 1). Ratings after reappraising negative content were more positive (mean negative reappraise rating = 4.25 ± 0.81 SD, Posterior Probability of Negative Regulate being greater than Negative View ratings (PP(Negative Regulate > Negative View Ratings)) > 0.9999) than after simply viewing negative scenes (mean negative view rating = 2.69 ± 0.54 SD; PP(Neutral View > Negative View Ratings) > 0.9999). Likewise, emotion ratings after reappraising positive stimuli (mean positive reappraise rating = 5.21 ± 0.9 SD; PP(Positive Regulate < Positive View Ratings) > 0.9999) were lower than after simply viewing positive content (mean positive view rating = 7.09 ± 0.67 SD; (PP(Positive View > Neutral View Ratings) > 0.9999)). Thus, participants were successful in regulating their emotional responses to the affective pictures when asked to do so.

**Figure 2.**
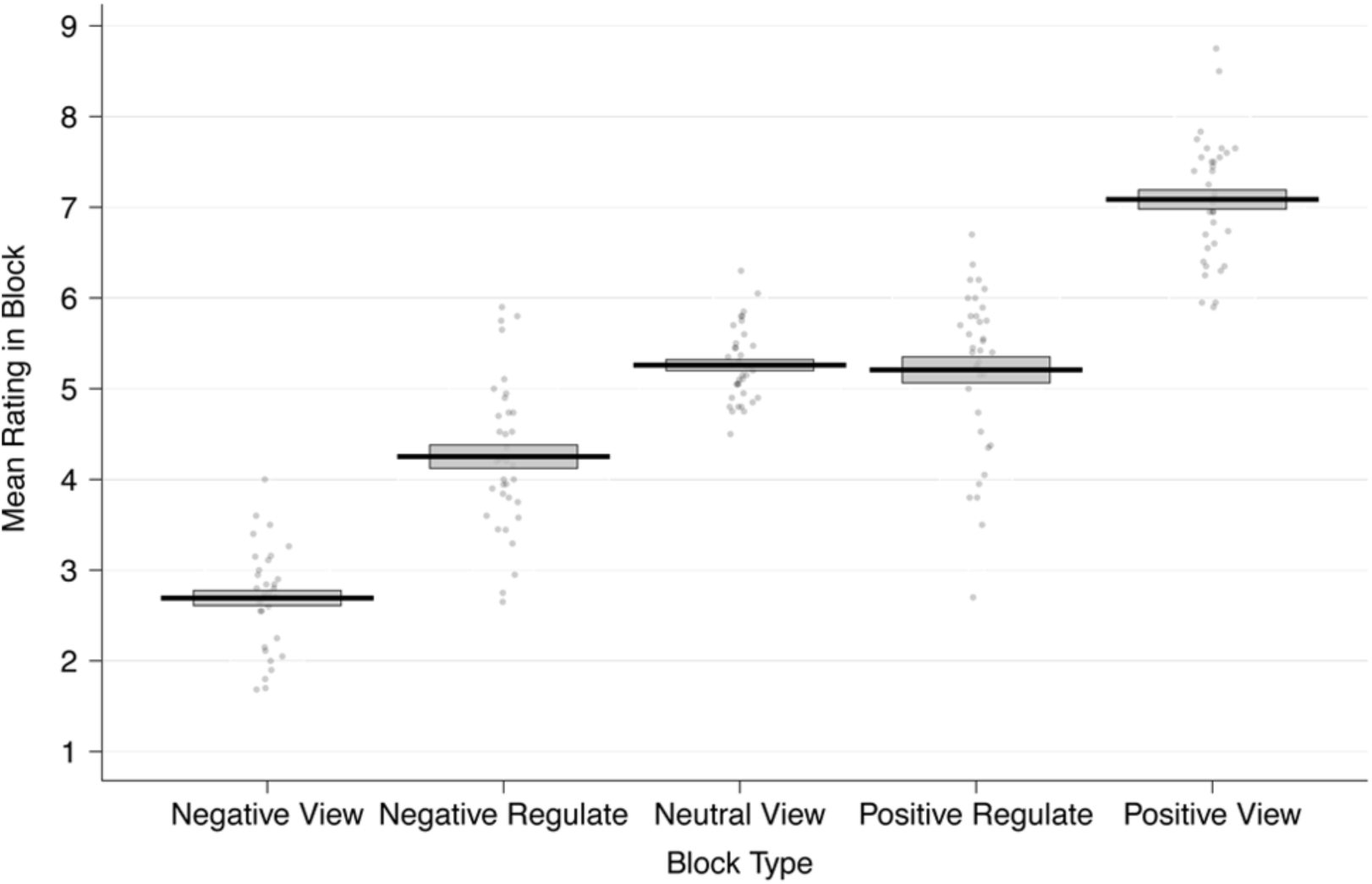
Emotion reappraisal behavior: Mean ratings made during the fMRI blocks by each participant. Ratings are aggregated over the negative view, negative regulate (reappraise), neutral view, positive regulate, and positive view blocks. Participants successfully reappraised negative images such that their emotions became more positive, and positive images such that their emotions became more negative. The black solid line represents the group mean and the gray box indicates the standard error of the mean. Each dot represents the mean ratings from one participant.

##### Dietary self-control task

In addition to the emotion regulation task, participants also completed a food choice task. The food choice task required subjects to make 100 decisions about whether or not they would eat the food item shown on the screen after the MRI scan. Participants knew that one of these trials would be selected at random and their choice on that trial implemented for real, meaning that they would have to eat the food item or go hungry for an additional 30 minutes. In analyzing the food choice behavior, we first examined the entire set of food choices using a mixed-effects logistic regression (see Eq. 2 and Table 2). This regression showed that, on average, participants considered both taste and health to a similar degree when choosing whether or not to eat the item shown on the screen (regression coefficient (coef.) taste = 1.47; coef. Health = 1.46). Consumption choices did not significantly differ as a function of task order, hunger levels, gender, BMI or restrained eating score (see Table 2).

**Table 2.**
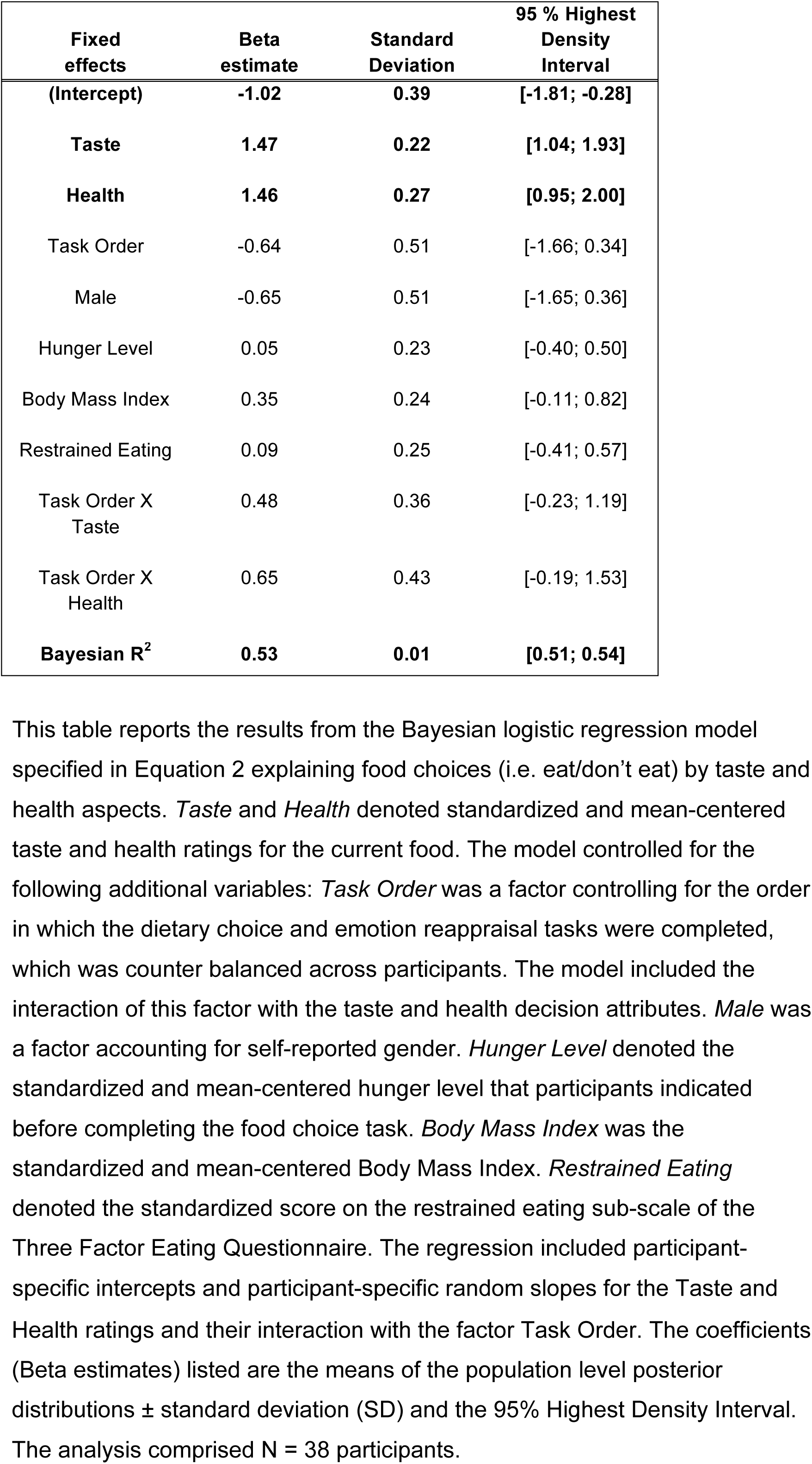
Basic food choice model.

Next, we focused specifically on food choices that represented a self-control challenge. These were trials in which the food was either palatable, but unhealthy, or healthy, but unpalatable according to the participants’ subjective ratings for healthiness and tastiness. Participants faced a self-control challenge on approximately 75 out of the 100 trials. The mean dietary self-control success level across all participants was 62 ± 27 SD %. This indicates that self-control success levels were high on average, but also that there was substantial variability across participants in dietary self-control. We also found that self-control success was achieved more often by refusing to eat tasty-unhealthy foods (Figure 3A). The mean self-control success level for refusing the tasty-unhealthy foods was 77% in our sample, whereas the mean success level for accepting unpalatable-healthy foods was only 19%.

**Figure 3.**
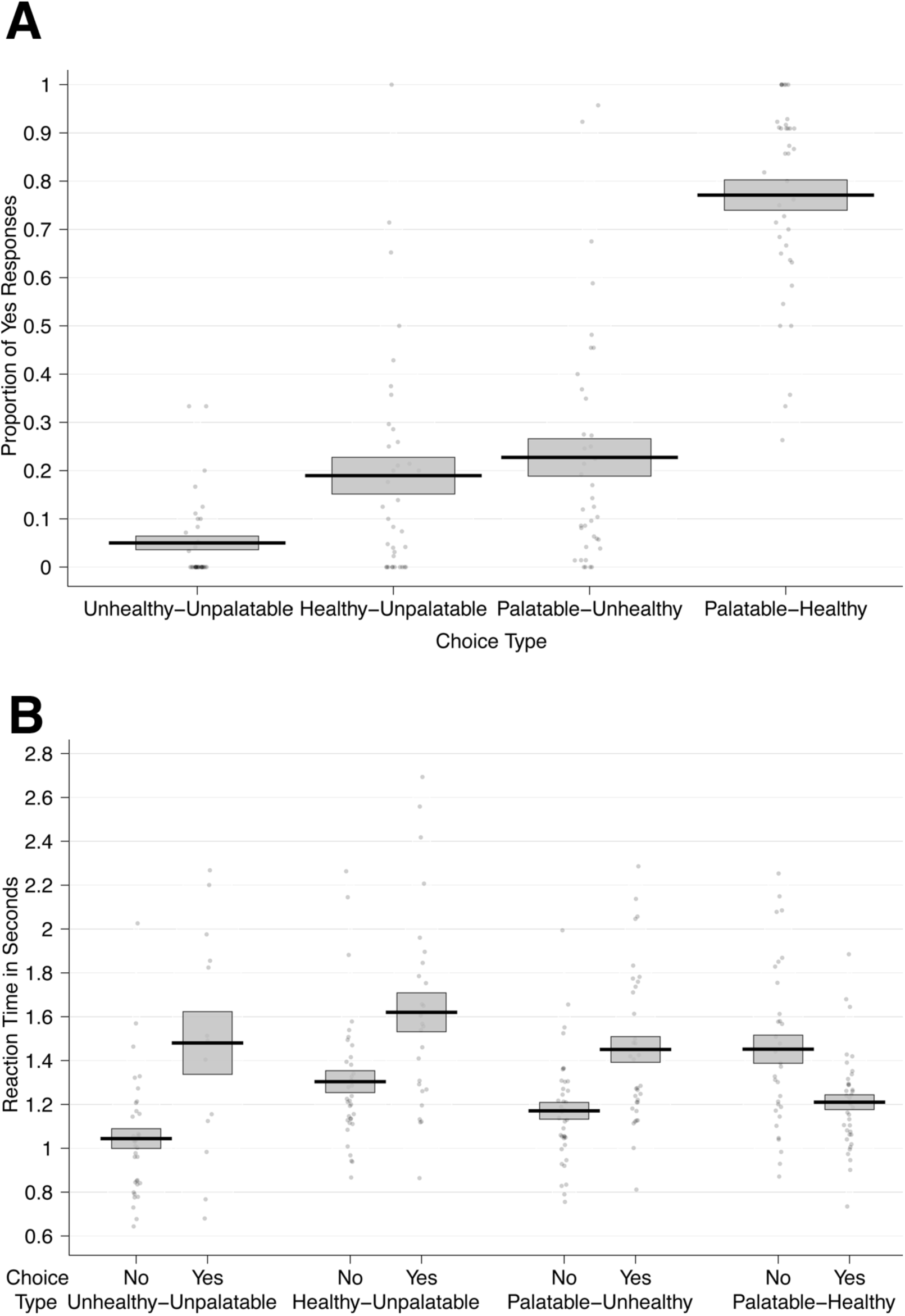
Dietary choice behavior: Panel **A** shows the proportion of “Yes” responses by choice category. Panel **B** shows the mean reaction times (RTs) over all participants for accepting and rejecting to eat foods from each of the four categories. In both panels, the black solid line represents the group mean and the gray box indicates the standard error of the mean. Each dot represents the proportion of “Yes” choices (A) or mean RT by choice (B) for one participant.

To test the influences of taste and health attributes and challenge type on self-control success, we performed a second mixed effects logistic regression (Eq. 3 and Table 3). Overall, the log odds of self-control success were lower for unpalatable-healthy foods (coef. = −1.89) compared to tasty-unhealthy foods. For tasty-unhealthy food, higher taste decreased the log odds of success (coef. = −1.40). Higher health ratings of the tasty-unhealthy foods also decreased the chances of refusing to eat them (coef. = −2.75), perhaps because choosing such a food with relatively higher healthiness might be perceived as a less serious failure. These results suggest that participants were not just habitually refusing tasty-unhealthy foods, because their choices were sensitive to both taste and health aspects in this type of challenge. For healthy-unpalatable food, relatively less bad-tasting foods increased the log odds of success (coef. = 2.67). However, healthiness had little influence on choice during healthy-unpalatable trials. Note that the total effect is equal to the Type x Health interaction coefficient (2.69) added to the baseline coefficient (−2.75). In other words, the significant influence of healthiness during unhealthy-palatable trials, which serve as the baseline in our regression, disappears (2.69 + −2.75 = −0.06) in healthy-unpalatable self-control challenges. The reduced influence of healthiness on these trials may be because the alternative of eating nothing at all for an extra 30 minutes is not viewed as an unhealthy outcome.

**Table 3.** Self-control success by taste and health attributes and challenge type.

| Fixed effects | Beta estimate | Standard Deviation | 95 % Highest Density Interval |
| --- | --- | --- | --- |
| (Intercept) | 1.28 | 0.29 | [0.73; 1.86] |
| Taste | -1.40 | 0.33 | [-2.04; -0.72] |
| Health | -2.75 | 0.44 | [-3.67; -1.94] |
| Type | -1.89 | 0.91 | [-3.77; -0.17] |
| Type X Taste | 2.67 | 0.58 | [1.63; 3.93] |
| Type X Health | 2.69 | 0.71 | [1.27; 4.09] |
| Bayesian R<sup>2</sup> | 0.62 | 0.01 | [0.61; 0.63] |

This table reports the results from the Bayesian logistic regression model specified in Equation 3 explaining dietary self-control success (coded as a binary variable: 1 = success / 0 = no success) by taste and health aspects as well as challenge type. *Taste* and *Health* denoted between-participant standardized and mean-centered taste and health ratings for the current food. *Type* was a factor accounting for the type of challenge (levels: 0 = high-taste/low-health, 1 = high-health/low-taste). The model included the interaction of this factor with the taste and health decision attributes. The regression included participant-specific intercepts and participant-specific random slopes for the taste and health ratings and their interaction with the challenge type. The coefficients (Beta estimates) listed are the means of the population level posterior distributions ± standard deviation (SD) and the 95% Highest Density Interval. The analysis comprised N = 38 participants.

We also examined reaction times (RT) for trials including healthy-unpalatable and palatable-unhealthy foods as well as trials in which taste and health attributes were aligned (Figure 3B). Notably, participants were faster to refuse eating the foods for all trial types, except when foods were both high in taste and health attributes (i.e. obvious eat decisions; see Figure 3B, Eq. 4 and Table 4). These RT results suggest that participants may have developed a bias toward refusing to eat the foods in this task. This can be seen in Table 4: The non-challenging trials (in which health and taste aspects were aligned) served as the baseline for this model. In these trials (approximately 25%), which were mostly high-taste/high-health foods, participants were significantly faster when they accepted to eat a food. Here, “Yes/Eat” responses are fast because the positive values of taste and healthiness quickly overcome the bias toward refusing the items (coef. = −0.07). In contrast, for both the high-taste/low-health (HTLH) and low-taste/high-health (LTHH) challenge trials (approximately 75% of all trials), participants were faster when they refused to eat the food compared to when they chose to eat it. This is consistent with an initial bias to refuse eating those conflicted foods, and overriding it requiring more time. Such a bias is potentially strategic for self-control because participants most often saw tasty-unhealthy foods in this task and, therefore, may have prepared in advance to decline eating them.

**Table 4.** Reaction time model for food choices.

| Fixed effects | Beta estimate | Standard Deviation | 95% Highest Density Interval |
| --- | --- | --- | --- |
| **Intercept** | **0.15** | **0.03** | **[0.09; 0.20]** |
| **Yes (to HTHH)** | **-0.07** | **0.03** | **[-0.12; -0.02]** |
| **HTLH** | **-0.05** | **0.02** | **[-0.09; -0.01]** |
| LTHH | 0.03 | 0.02 | [-0.02; 0.07] |
| Stakes | 0.00 | 0.01 | [-0.03; 0.03] |
| **Difficulty** | **0.10** | **0.03** | **[0.05; 0.15]** |
| **Yes X HTLH** | **0.24** | **0.05** | **[0.15; 0.33]** |
| **Yes X LTHH** | **0.26** | **0.05** | **[0.17; 0.36]** |
| Bayesian $R^2$ | 0.27 | 0.01 | [0.25; 0.29] |

This table reports the results from the Bayesian regression model of reaction times for food choices specified in Eq. (4). Reaction times were transformed using the natural logarithm. The variable *Yes* was coded with a value of 1 if participants chose to eat the depicted item and 0 otherwise. Trial type was coded as a factor with 3 categories (non-challenging high-taste/high-health (HTHH) and low-taste/low-health (LTLH) trials as the reference category, and high-taste/low-health (*HTLH*) trials and low-taste/high-health (*LTHH*) trials as indicator variables). The variable *Stakes* was calculated for each trial as described in Eq. (5) and the variable *Difficulty* was calculated for each trial as described in Eq. (6). The regression included participant-specific intercepts and participant-specific random slopes for all regressors and interaction terms. The coefficients (Beta estimates) listed are the means of the population level posterior distributions ± standard deviation (SD) and the 95% Highest Density Interval. The analysis comprised $N = 38$ participants.

Further corroboration for the existence of this initial bias comes from drift-diffusion modeling results that show that in the current study, participants have a starting-point bias (approximately 1/3) toward refusing the foods across all trials (see Supplementary Methods, Results, and Table S2). Response times generated using this starting-point bias and the other best fitting DDM parameters (Supplementary Figure S2) reproduce the RT pattern seen in Figure 3. Specifically, the Yes/Eat decisions are slower than No/Don’t eat decisions in all cases, except when Yes is the obvious decision because both palatability and healthiness are high.

Notably, the reaction time patterns for the low-taste/high-health foods speak against a default strategy of choosing healthy: Pondering whether to eat the food or not in such choices slowed down participants substantially (coef. = 0.26). If the participants just followed an often-practiced health habit from their daily life, this should not be the case. If they followed such a habit, then choosing healthy foods should come to them naturally and quickly. The slowdown suggests participants perceived a challenge and /or had to overcome a bias to refuse eating the foods. Thus, the overall pattern of results is consistent with the idea that participants may have formed a bias to refuse eating the foods.

#### Testing behavioral associations between tasks

Next, in order to address our questions about the potential link between emotional reappraisal and dietary self-control at the behavioral level, we tested for an association between the self-reported reappraisal and dietary self-control success scores. However, we did not observe a significant correlation between overall dietary self-control success level and emotional reappraisal success (Bayesian rank correlation rho = −0.023, 95% Highest Density Interval (HDI) = [-0.368; 0.306], posterior probability of rho greater than zero (PP rho > 0) = 0.450). For completeness, we also ran separate tests for reappraisal success in the positive (rho = 0.136, 95% HDI = [-0.199; 0.472], (PP rho > 0) = 0.781) and negative valence domains (rho = −0.175, 95% HDI = [-0.499; 0.156], (PP rho > 0) = 0.159), but these did not show significant correlations with the overall dietary self-control success level either.

### fMRI

#### Testing for previously observed patterns of BOLD activity within each task

Before testing for associations between dietary self-control and emotion regulation at the neural level, we first checked if the patterns of neural activity within each paradigm were consistent with previous findings from emotion reappraisal and dietary choice studies.

##### Reappraisal task

Our findings from the reappraisal paradigm were consistent with past fMRI studies examining the neural correlates of reappraising emotional scenes. The contrast of Reappraisal Success > View across both positive and negative valence showed several regions noted in previous work (Gross, 1998; Ochsner and Gross, 2005; Wager et al., 2008; Ochsner et al., 2012; Buhle et al., 2014; Etkin et al., 2015; Morawetz et al., 2017a) such as medial temporal gyrus, SMA, caudate, putamen, insula, vlPFC and dlPFC were more active when reappraising emotional scenes compared to viewing them and reacting naturally (GLM-ER; Figure 4, Table 5). Note that the contrasts Reappraisal Success > View and Reappraisal > View were very similar in this sample. This in not surprising because participants rarely failed to reappraise the image content (see Supplementary Figure S3 and Table S3). In line with the behavioral finding that participants succeeded in reappraising both valences, positive and negative emotion reappraisal success did not significantly differ in terms of BOLD activity (Negative Reappraisal Success > Positive Reappraisal Success: all p-values > 0.28, whole-brain family-wise error corrected; Positive Reappraisal Success > Negative Reappraisal Success: all p-values > 0.29 whole-brain corrected).

**Figure 4.**
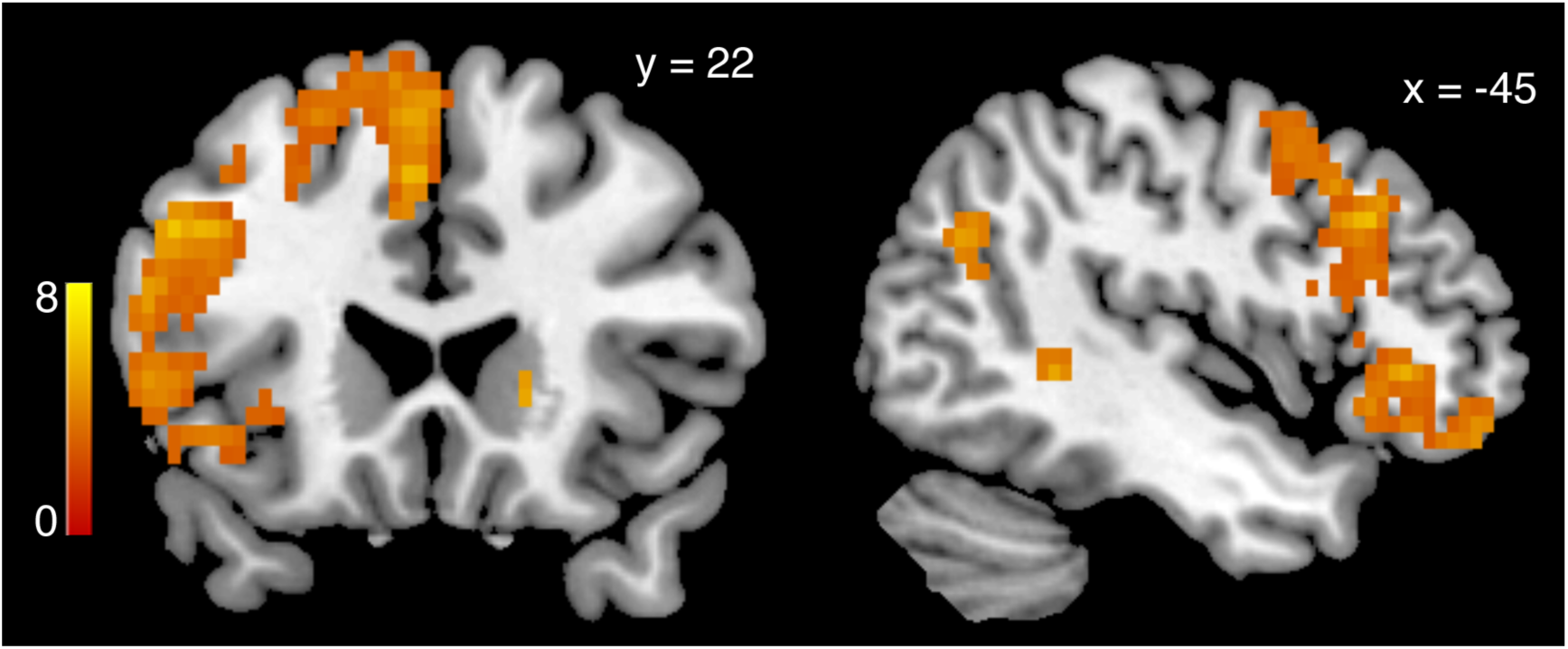
Successfully Reappraising > Viewing emotional content: Collapsed over both positive and negative stimuli, BOLD activity in GLM-ER was greater in a widespread set of brain regions when successfully reappraising the content of emotional pictures in order to dampen the elicited emotions, compared to viewing the stimuli without altering the elicited feeling (p < 0.05, whole-brain corrected, derived from 5000 permutations of the data). The heat map represents T-statistics on a scale from 0 to 8 to keep the scale consistent across all subsequent figures.

**Table 5.**
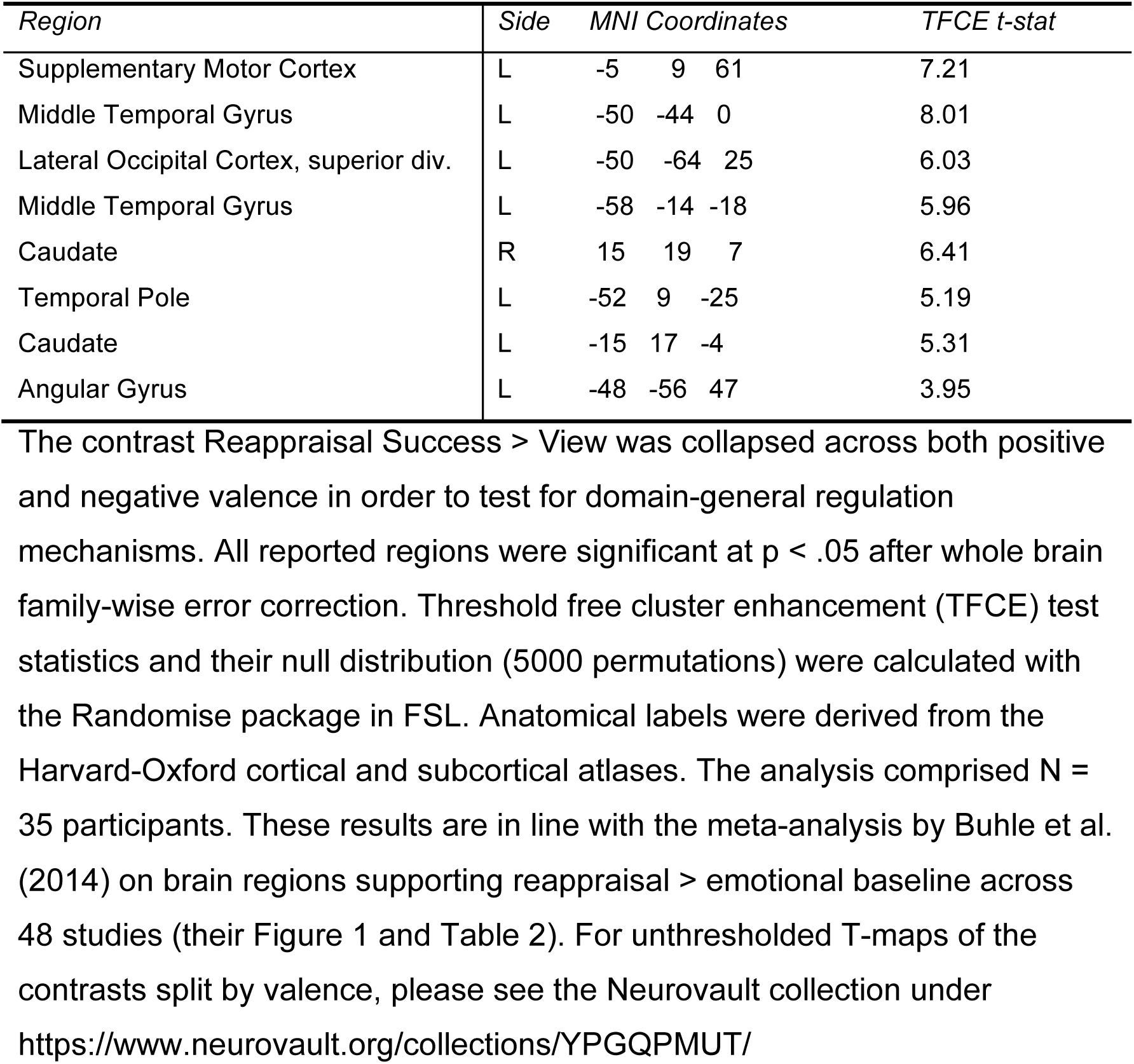
Reappraisal Success > View in GLM-ER.

##### Dietary self-control task

In the food choice task, some of our analyses were consistent with previous reports, but in other cases there were notable differences. Using GLM-FC, we found BOLD activity scaling with subjective food value in a set of brain regions typically associated with value-based choices during tests of self-control (Hare et al., 2009; Hare et al., 2011; Enax et al., 2015; Maier et al., 2015; Spetter et al., 2017; van Meer et al., 2017) and more generally (i.e. without explicit self-control) (Bartra et al., 2013; Clithero and Rangel, 2013). These included the medial prefrontal and posterior cingulate cortices (Figure 5A, Table 6; p = 0.01, whole-brain corrected). A separate GLM (GLM-TH) that replaced the subjective food values with the individual taste and healthiness ratings showed that overlapping regions also represented healthiness (Figure 5B, Table 7; p < 0.0001) and tastiness (Figure 5C, Table 8; p = 0.02) attributes. Figure 5D depicts the overlap between the regions that significantly encoded subjective food value, taste and health (conjunction threshold = p < 0.05, whole-brain corrected for family-wise error).

**Figure 5.**
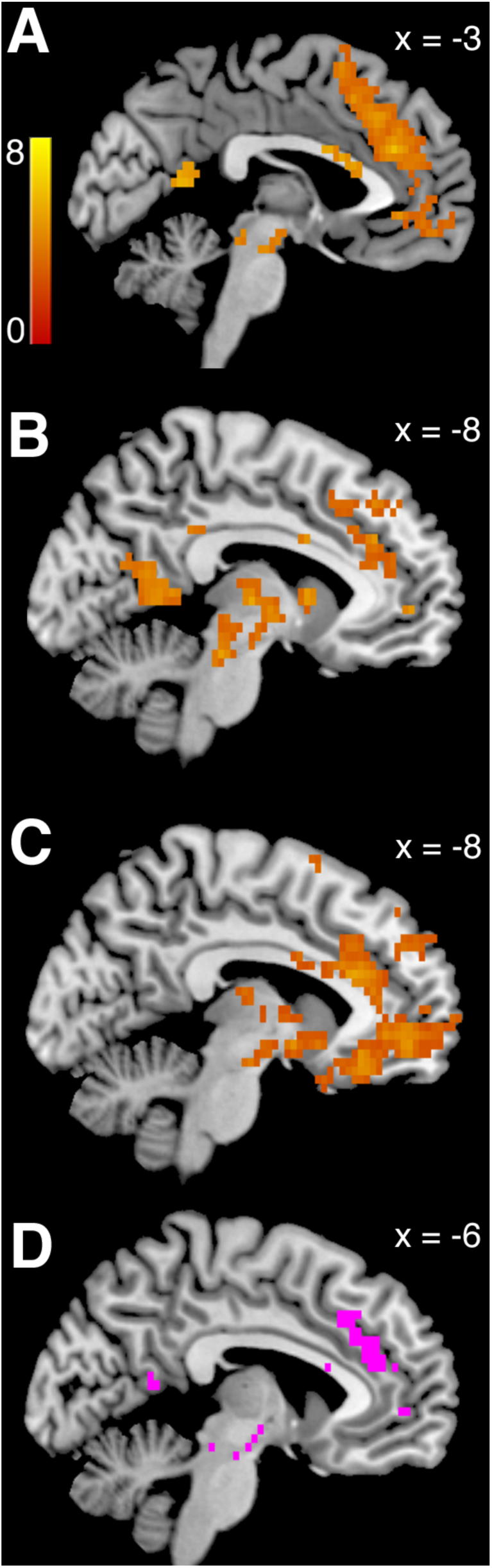
Neural activity at the time of food choice: **A)** BOLD activity increased with higher subjective food value in a set of regions associated with value-based choice in GLM-FV. Panel **B)** depicts regions that increased their BOLD activity with higher health ratings, and panel **C)** regions that increased their activity with higher taste ratings of the presented foods (both from GLM-TH). All results in panels A)-C) were significant at the threshold of p < 0.05, whole-brain corrected. The heat map represents T-statistics derived from 5000 permutations of the data. Panel **D)** depicts in pink the overlap of areas that significantly encoded the three-way conjunction of subjective food value, taste and health attributes. The conjunction threshold was p < 0.05, whole-brain corrected for family-wise error.

**Table 6.** Subjective food value representations in GLM-FV.

| *Region* | *Side* | *MNI Coordinates* |  |  | *TFCE t-stat* |
| --- | --- | --- | --- | --- | --- |
| Paracingulate Gyrus | L | -3 | 37 | 25 | 5.99 |
| Brain Stem | R | 10 | -26 | -14 | 5.79 |
| Anterior Cingulate Gyrus | R | 0 | 14 | 22 | 5.43 |
| Precuneus | L | -3 | -56 | 11 | 5.59 |
| Orbital Frontal Cortex | L | -28 | 32 | -14 | 4.36 |
| Insular Cortex / Orbital Frontal Cortex | L | -30 | 9 | -14 | 5.09 |
| White Matter | R | 10 | 17 | 22 | 4.36 |
| Medial Frontal Cortex | L | -10 | 34 | -18 | 4.65 |
| Orbital Frontal Cortex | L | -25 | 24 | -22 | 4.48 |
| Amygdala | R | 13 | -6 | -11 | 5.31 |

All reported regions were significant at $p < .05$ after whole brain family-wise error correction. Threshold free cluster enhancement (TFCE) test statistics and their null distribution (5000 permutations) were calculated with the Randomise package in FSL. Anatomical labels were derived from the Harvard-Oxford cortical and subcortical atlases. The analysis comprised $N = 37$ participants.

**Table 7.** Health value representations in GLM-TH.

| *Region* | *Side* | *MNI Coordinates* |  |  | *TFCE t-stat* |
| --- | --- | --- | --- | --- | --- |
| Superior Frontal Gyrus | L | -13 | 37 | 47 | 6.26 |
| Frontal Pole | L | -45 | 42 | 14 | 5.63 |
| Caudate | L | -10 | 9 | 7 | 5.32 |
| Precuneus | L | -3 | -56 | 11 | 6.13 |
| Posterior Cingulate Gyrus | L | -5 | -36 | 32 | 5.45 |
| Caudate | R | 13 | 9 | 11 | 5.73 |
| Anterior Cingulate Gyrus | L | -3 | 7 | 25 | 5.28 |
| White Matter | R | 10 | 17 | 22 | 5.66 |
| Anterior Cingulate Gyrus | L | -3 | 22 | 18 | 4.25 |
| Superior Lateral Occipital Cortex | L | -35 | -66 | 47 | 5.7 |
| Paracingulate Gyrus | L | -8 | 52 | 0 | 4.77 |
| Anterior Cingulate Gyrus | L | -3 | 14 | 22 | 4.22 |

This table lists regions significantly correlated with the parametric modulator for health ratings in GLM-TH. All reported regions were significant at $p < .05$ after whole brain family-wise error correction. Threshold free cluster enhancement (TFCE) test statistics and their null distribution (5000 permutations) were calculated with the Randomise package in FSL. Anatomical labels were derived from the Harvard-Oxford cortical and subcortical atlases. The analysis comprised $N = 37$ participants.

**Table 8.** Taste value representations in GLM-TH.

| *Region* | *Side* | *MNI Coordinates* |  |  | *TFCE t-stat* |
| --- | --- | --- | --- | --- | --- |
| Orbital Frontal Cortex | R | 25 | 24 | -18 | 6.02 |
| Frontal Pole | R | 18 | 47 | 36 | 4.39 |
| Supplementary Motor Cortex | L | -3 | 7 | 65 | 4.13 |
| Posterior Cingulate Gyrus / Precuneus | L | -3 | -51 | 14 | 4.7 |
| Anterior Parahippocampal Gyrus | R | 20 | -19 | -29 | 4.03 |
| Superior Frontal Gyrus | L | -15 | 27 | 54 | 3.9 |

This table lists regions significantly correlated with the parametric modulator for taste ratings in GLM-TH. All reported regions were significant at $p < .05$ after whole brain family-wise error correction. Threshold free cluster enhancement (TFCE) test statistics and their null distribution (5000 permutations) were calculated with the Randomise package in FSL. Anatomical labels were derived from the Harvard-Oxford cortical and subcortical atlases. The analysis comprised $N = 37$ participants.

Previous studies have reported that self-control success is associated with greater activity in the dorsolateral prefrontal and occipital cortex (Hare et al., 2009; Christakou et al., 2011; Crockett et al., 2013; Harris et al., 2013; Drobetz et al., 2014; Schonberg et al., 2014; Decker et al., 2015; Luerssen et al., 2015; Maier et al., 2015; Hill et al., 2017; Spetter et al., 2017; Baumeister et al., 2018; Bertsch et al., 2018; Jimura et al., 2018; Lee et al., 2018; Schmidt et al., 2018; Shahbabaie et al., 2018; Sheffer et al., 2018). However, in the current dataset, we did not find in GLM-SCS greater activity in the dlPFC, or any other brain regions, on Successful Self-control trials compared to Self-control Failures (all p-values > 0.21, whole-brain corrected; all p > 0.65 small-volume corrected in left lateral PFC). Similarly, the contrast for Self-control Success > No Challenge yielded no significant difference in the prefrontal cortex (all p-values > 0.37, whole-brain corrected; all p > 0.77, small-volume corrected in left lateral PFC). Individual differences in the overall dietary self-control success level did not correlate with activity in any prefrontal regions, but we did find that greater activity within the left lingual and fusiform gyri (Table 9; p = 0.01, whole-brain corrected) during Self-control Success vs. No-Challenge trials was positively correlated with the individual overall dietary self-control success levels. These results linking self-control to activity in regions involved in visual and object processing are consistent with both the speculations about early filtering of visual attention as a mechanism to facilitate self-control in Harris et al. (2013) and the pattern of fast refusals for unhealthy foods observed in the current participants’ behavior.

**Table 9.** Correlation of individual differences in overall dietary self-control success level with the Self-Control Success > No Challenge contrast from GLM-SCS.

| *Region* | *Side* | *MNI Coordinates* | *TFCE t-stat* |
| --- | --- | --- | --- |
| Occipital fusiform gyrus | L | -25 -86 -11 | 5.65 |

The reported region was significant at $p = 0.01$ after whole brain family-wise error correction. Threshold free cluster enhancement (TFCE) test statistics and their null distribution (5000 permutations) were calculated with the Randomise package in FSL. Anatomical labels were derived from the Harvard-Oxford cortical and subcortical atlases. The analysis comprised $N = 37$ participants.

#### BOLD activity during emotion reappraisal is associated with dietary self-control success

Next, we tested the hypotheses that neural activity patterns during successful reappraisal would be related to individual differences in dietary self-control success or vice versa. We computed a between-subjects regression relating individual differences in overall dietary self-control success levels to voxel-wise differences in the Reappraisal Success > View contrast. This analysis revealed that participants whose BOLD signal changed more strongly when successfully reappraising compared to viewing emotional content were also overall better at dietary self-control (Figure 6A, Table 10; p < 0.025 whole-brain corrected). We observe this correlation in areas that represent the value assigned to foods during the food choice task (Figure 6B). These food values are based on the relative importance of the palatability and health attributes of the foods and are closely associated with the decision to eat or forego the food on every trial (see Table 2). Thus, the areas, in which activity during the emotion regulation task is linked to a participant’s average level of dietary self-control, also correlate with trial-wise value computations that support dietary decisions in the food choice task itself.

**Figure 6.**
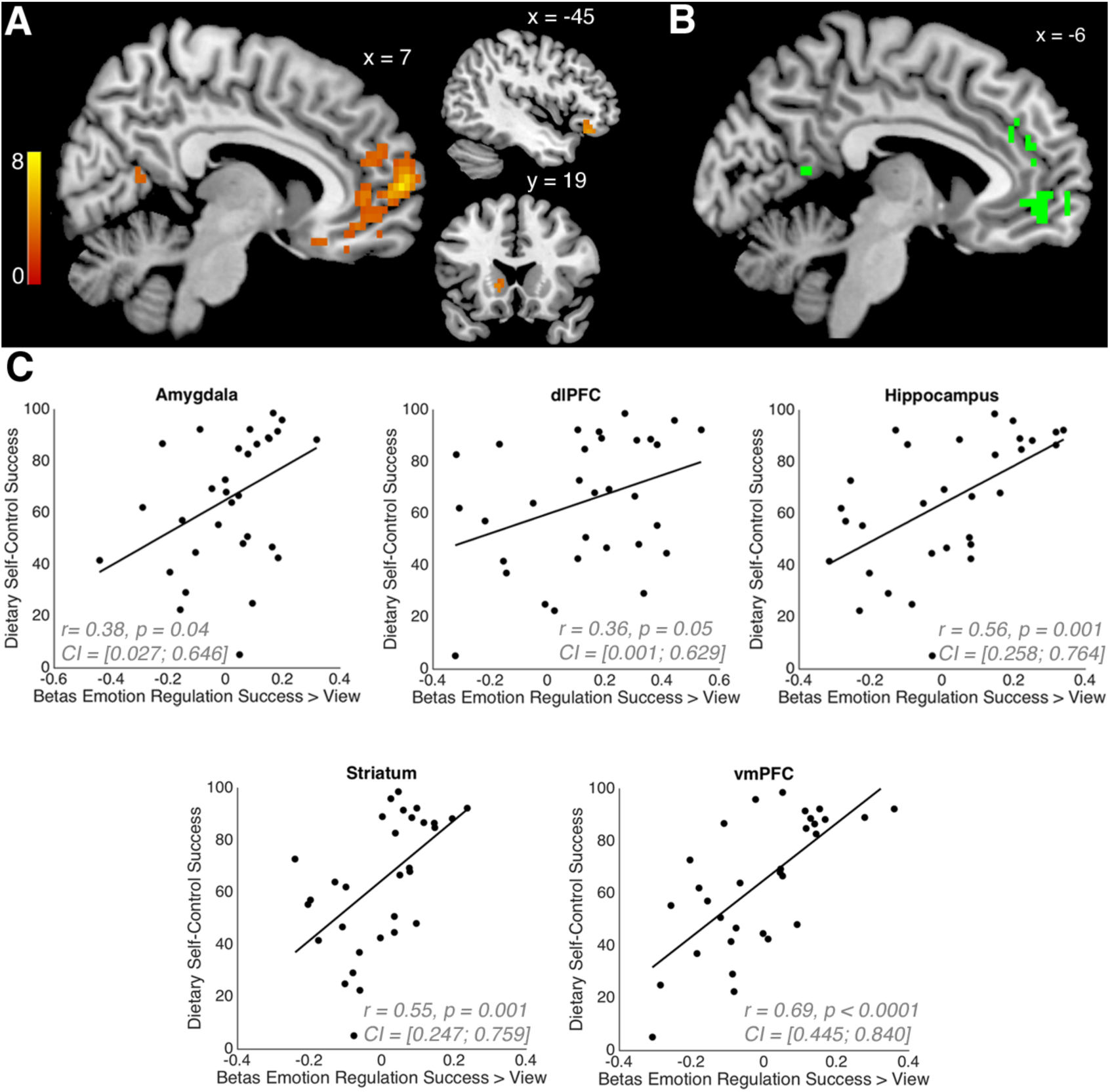
Emotion reappraisal and dietary self-control link: Panel **A)** shows the results from a between-subjects regression relating individual differences in the overall dietary self-control success level to voxel-wise differences in BOLD activity during emotion regulation. Activation when successfully reappraising compared to viewing emotional content (GLM-ER) was higher in participants with better dietary self-control (p < 0.025, whole-brain corrected, T-statistics derived from 5000 permutations of the data). This suggests that participants whose neural activity changed more strongly during reappraisal of positive and negative stimuli were also the ones who were better at modulating their dietary decisions to refuse eating tasty-unhealthy foods or increase eating healthy-untasty foods. Panel **B)** shows the conjunction of the contrasts “Subjective Food Value” from GLM-FV (panel A in Figure 5) and the between-subjects regression contrast from GLM-ER (panel A in Figure 6). The overlap suggests that regions in the medial prefrontal and posterior cingulate cortices may be involved in modifying the subjective valuation of emotional stimuli and the computation of subjective food values that factor in both health and taste attributes. The scatter plots in panel **C)** illustrate the strength of the relationship between emotion reappraisal and dietary self-control success shown in panel A. We performed region-of-interest (ROI) analyses in 5 regions that have previously been associated with reappraisal and decision-making in order to assess how strongly the BOLD activity change in Reappraisal > View conditions was related to the overall dietary self-control success level. The statistics in each plot give Pearson’s rho (r) and its 95% Confidence Interval (CI) as well as the p-value (two-sided test) for the correlations between the mean BOLD activity for the contrast Reappraisal Success > View in each region and the overall dietary self-control success level (in percent).

**Table 10.** Reappraisal Success > View (in GLM-ER) correlates with overall dietary self-control success levels.

| *Region* | *Side* | *MNI Coordinates* | *TFCE t-stat* |
| --- | --- | --- | --- |
| Frontal Pole | R | 8 62 7 | 7.33 |
| Precuneus | L | -3 -56 7 | 4.25 |
| Temporal Occipital Fusiform Cortex | R | 43 -61 -25 | 4.35 |
| Cerebellum | L | -20 -86 -32 | 4.91 |
| Caudate | L | -13 19 0 | 4.14 |
| Posterior Parahippocampal Gyrus | L | -18 -26 -11 | 4.48 |
| Orbital Frontal Cortex | L | -45 32 -14 | 4.96 |
| Posterior Cingulate Gyrus | L | -3 -34 7 | 4.03 |
| Lingual Gyrus / Occipital Pole | L | -3 -91 -18 | 4.05 |
| Frontal Pole | L | -25 42 7 | 3.73 |
| Cerebellum | R | 28 -84 -32 | 4.74 |
| Lingual Gyrus | R | 15 -64 -11 | 5.1 |
| Posterior Parahippocampal Gyrus | R | 23 -34 -18 | 3.86 |
| Temporal (Occipital) Fusiform Cortex | R | 33 -39 -25 | 3.7 |
| Cerebellum | R | 3 -89 -29 | 3.68 |
| Precuneus | R | 13 -56 14 | 3.3 |
| Corpus Callosum | L | -5 27 7 | 3.48 |
| Planum Polare | L | -43 2 -18 | 5.04 |
| Cerebellum | R | 30 -59 -25 | 3.13 |
| Cerebellum | R | 35 -59 -58 | 4.37 |

All reported regions were significant at $p < .025$ after whole brain family-wise error correction. We Bonferroni-corrected for multiple comparisons because two separate analyses were required to obtain this result (assessment of the overall dietary self-control success level, and assessment of neural reappraisal correlates). Threshold free cluster enhancement (TFCE) test statistics and their null distribution (5000 permutations) were calculated with the Randomise package in FSL. Anatomical labels were derived from the Harvard-Oxford cortical and subcortical atlases. The analysis comprised $N = 31$ participants.

We additionally conducted a-priori region-of-interest (ROI) analyses in regions that have previously been associated with reappraisal and self-control decisions. We computed Pearson’s correlation coefficient between the overall dietary self-control success level and Reappraisal Success > View BOLD activity from the following regions (bilateral, anatomically defined based on the Harvard-Oxford Atlas): amygdala, hippocampus, striatum (nucleus accumbens, caudate and putamen) and vmPFC as well as left dlPFC (from the union of voxels associated with self-control in Hare et al. (2009) or Maier et al. (2015)). We identified positive correlations between the contrast of Reappraisal Success > View and the overall dietary self-control success level in vmPFC, hippocampus and striatum (Bonferroni-corrected for multiple comparisons: all p-values < 0.01; see Figure 6C).

The complementary test for whether BOLD activity differences in the dietary Self-control Success > No Challenge contrast from GLM-SCS were linked to overall emotion reappraisal success scores did not yield a significant correlation in any regions (all p-values > 0.35 after whole-brain correction). Potential reasons for this asymmetry in the relationship between BOLD activity and regulation success across domains are considered in the Discussion section.

#### Prefrontal cortex BOLD signals correlate with dietary self-control stakes

Both the unexpected tendency to quickly refuse palatable and unpalatable food items and the lack of a significant relationship between PFC activity and dietary self-control prompted us to conduct additional exploratory analyses on the fMRI data from the food choice task. One question we had was if participants were tracking the healthiness and tastiness attributes at stake on each trial with regard to the need for self-control even though they seemed to have a bias toward declining to eat the food items. Given the previous findings implicating left PFC in dietary self-control cited above, we initially searched there. We found that a measure of the objective self-control stakes (defined as |HR| + |TR| on challenge trials, see GLM-ST) was correlated with BOLD signals in left Inferior Frontal Gyrus (IFG) during dietary decisions (Figure 7C, blue areas; Table 11, p = 0.01, svc within left prefrontal cortex). A post-hoc comparison of the average coefficients for taste and healthiness stakes (i.e. |TR| or |HR|) within this functional ROI indicated that the left IFG region represented both attributes, rather than tracking only one or the other. Lastly, a whole-brain analysis revealed a trend for a bilateral activation of the IFG (with additional activation in the right IFG: peak MNI coordinate = [55 29 0], max T = 5.49, p = 0.06 whole-brain corrected), suggesting that this pattern is not strictly lateralized. Thus, this initial set of exploratory analyses indicated that the BOLD activity in the IFG is correlated with the size of the stakes for self-control challenges.

**Figure 7.**
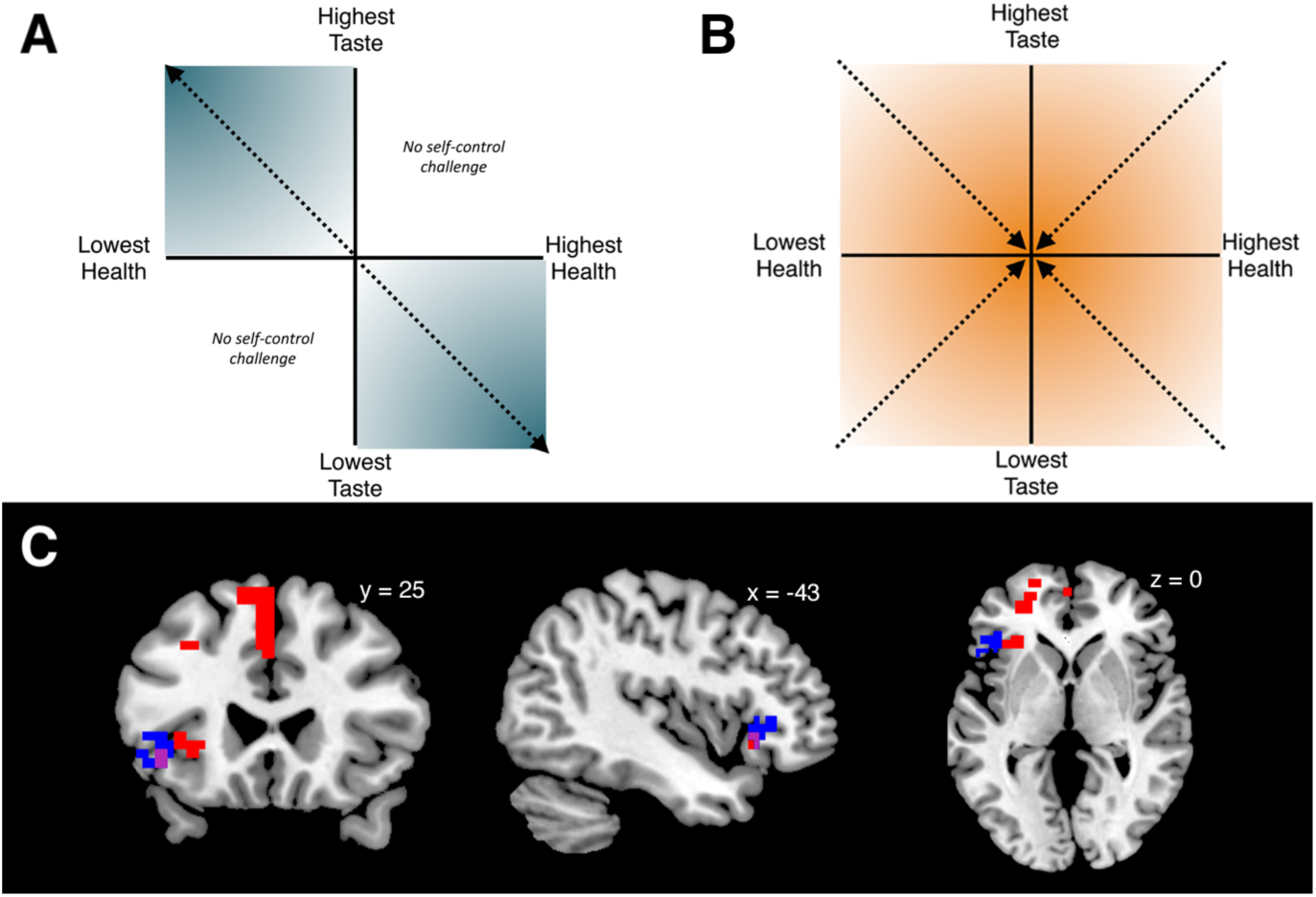
Self-control stakes: The sketch in panel **A)** explains the intuition for quantifying what is at stake in self-control. In the dietary self-control paradigm, any food can be categorized in one of four combinations of taste and health: tasty-healthy foods (upper right quadrant) and foods that are neither tasty nor healthy (lower left quadrant) present no challenge to self-control. When taste and health are not aligned, as foods become tastier and less healthy, the need for self-control increases (upper left quadrant). The same is true for the lower right quadrant as foods become healthier and a higher desire to eat tasty needs to be overcome. The intensifying shading illustrates how both aspects become more important the farther from zero (the middle of the neutral zone of the rating scale) participants rated each aspect. Thus adding up the distance from zero for taste and health (|tr| + |hr|) determines the self-control stakes. Note that the self-control stakes value is defined to be zero throughout the entire upper right and lower left quadrants. Panel **B)** illustrates decision conflict or choice difficulty. In contrast to the stakes of self-control that increase with higher distance from zero, choices become more difficult when the food value approaches zero, which means the options of eating the food or nothing are very similar. Furthermore, choice difficulty can have non-zero values in all four quadrant, unlike self-control stakes. Panel **C)** shows regions tracking the self-control stakes from GLM-ST: BOLD activity in the lateral prefrontal cortex (PFC) increased with higher stakes or importance of self-control (p < 0.05, small-volume corrected within left lateral PFC, T-statistics derived from 5000 permutations of the data). The voxels in blue or purple indicate the results from the current sample. To further test the relationship between stakes level and BOLD activity in these voxels, we conducted an ROI analysis using the functionally defined cluster from the current sample (i.e. blue and purple voxels) as a mask. We tested whether BOLD activity in a prior, independent study (Maier et al., 2015) showed the same association and found that BOLD signals in that sample also positively scaled with the trial-wise stakes level in these voxels (p = 0.04, T = 1.7472, df = 50). The results of a whole brain analysis for the stakes contrast in the Maier et al. (2015) sample are shown in red. Voxels in purple represent the conjunction of the contrasts from both datasets. In both datasets, the need for self-control was tracked by voxels in the left inferior frontal gyrus, while the larger sample from Maier et al. (2015) also identifies additional voxels in medial and dorsolateral prefrontal cortex.

**Table 11.** . Regions tracking the self-control stakes parametric modulator from GLM-ST in the current study.

| *Region* | *Side* | *MNI Coordinates* | *TFCE t-stat* |
| --- | --- | --- | --- |
| Frontal operculum / Inferior Frontal Gyrus | L | -45 29 0 | 5.23 |

The reported region was significant at $p = 0.01$ after small-volume family-wise error correction in a mask of the left prefrontal cortex. Threshold free cluster enhancement (TFCE) test statistics and their null distribution (5000 permutations) were calculated with the Randomise package in FSL. Anatomical labels were derived from the Harvard-Oxford cortical and subcortical atlases. The analysis comprised $N = 37$ participants.

**Table 12.** Regions tracking self-control stakes (i.e. the parametric modulator for stakes in GLM-ST) during self-control challenges in the Maier et al. (2015) dataset.

| **A. Small-volume corrected in left lateral PFC** |  |  |  |  |  |
| --- | --- | --- | --- | --- | --- |
| *Region* | *Side* | *MNI Coordinates* |  |  | *TFCE t-stat* |
| Superior Frontal Gyrus / Paracingulate Gyrus | L | -2 | 38 | 40 | 5.43 |
| Orbital Frontal Cortex | L | -30 | 28 | -4 | 4.48 |
| Middle Frontal Gyrus | L | -32 | 23 | 40 | 4.5 |
| Middle Frontal Gyrus | L | -40 | 16 | 43 | 4.14 |
| Orbital Frontal Cortex | L | -40 | 31 | -10 | 4.1 |
| Frontal Operculum | L | -37 | 26 | 3 | 3.77 |
| Orbital Frontal Cortex | L | -45 | 23 | -7 | 4.3 |
| **B. Whole-brain corrected** |  |  |  |  |  |
| Paracingulate Gyrus | R | 6 | 36 | 37 | 6.38 |
| Precuneus | R | 13 | -65 | 34 | 5.35 |
| Middle Frontal Gyrus | R | 41 | 18 | 46 | 4.29 |
| Middle Frontal Gyrus | R | 36 | 26 | 46 | 5.05 |
| Superior Lateral Occipital Cortex | R | 53 | -60 | 43 | 5.31 |
| Frontal Operculum | R | 43 | 23 | -4 | 4.39 |
| Angular Gyrus | R | 51 | -55 | 34 | 4.96 |
| Insula | L | -30 | 21 | -7 | 5.01 |

**A)** All reported regions were significant at $p < .05$ after small-volume correction for family-wise error in an anatomical mask of the left lateral prefrontal cortex. **B)** All reported regions were significant at $p < .05$ after whole-brain correction for family-wise error. Threshold free cluster enhancement (TFCE) test statistics and their null distribution (5000 permutations) were calculated with the Randomise package in FSL. Anatomical labels were derived from the Harvard-Oxford cortical and subcortical atlases. The analyses in both A) and B) comprised all 51 participants of the study.

In order to test whether the correlation between IFG activity and self-control stakes could be replicated in another independent dataset, we went back to the food choice and fMRI dataset first reported in Maier et al. (2015). We estimated the same BOLD GLM (i.e. GLM-ST) on these data, and found that indeed, the BOLD signal during challenging trials tracked the stakes in this independent dataset as well. Using the same left lateral prefrontal cortex mask as a small-volume search space, we observed activity in a region of the IFG that overlapped with the results from the current study (Figure 7C, purple areas represent overlap between current and prior datasets), as well as the medial PFC (Figure 7C, red areas; Table 12A; p < 0.0001 svc within left prefrontal cortex). A whole-brain analysis of the Maier et al. (2015) dataset revealed activity tracking the stakes in a large set of bilateral prefrontal voxels in the medial frontal gyrus, Brodmann areas 9 and 10, anterior cingulate, Brodmann areas 8 and 32 as well as the supplementary motor area (Table 12B). These results show that BOLD activity in prefrontal cortex correlates with the combined taste and healthiness outcome at stake during dietary self-control challenges in two independent datasets. We also conducted an ROI analysis within the region identified in the current dataset (Figure 7C, blue areas) and found that the stakes were also represented there in the dataset of Maier et al. 2015 (one-sample t-test, one-tailed hypothesis: betas are bigger than zero; p = 0.04, T = 1.7472, df = 50).

## Discussion

We demonstrated an association between BOLD activity during the successful reappraisal of emotional stimuli and the level of overall dietary self-control shown in a separate food choice task. Specifically, greater increase in BOLD signals in a distributed set of cortical and subcortical regions during successful emotional reappraisal was associated with better dietary self-control. Notably, many of the regions that showed this cross-domain correlation were more active for successful relative to failed reappraisal trials within our current and in previous emotion regulation experiments (see Buhle et al. (2014) for review). Together these results are consistent with the idea that neural processes related to the reappraisal of emotional stimuli may also facilitate dietary self-control.

### Association between the tasks

Despite the seemingly straightforward answer to one of the questions motivating our experiments, our findings also contain surprises that raised intriguing questions and prompted us to conduct further analyses. For example, the relationship between BOLD activity and regulation success across tasks was not symmetric. We didn’t find a significant relationship between BOLD activity during dietary self-control and self-reported reappraisal success. This may mean that stimulus reappraisal is one means of facilitating dietary self-control, but that the neural processes mediating dietary self-control are not directly relevant to stimulus reappraisal.

However, there are several other plausible explanations for this asymmetric relationship. The lack of correlation between food choice BOLD activity and reappraisal success may also be due to individual differences in how the affective ratings are subjectively reported. Recall that we have only subjective self-reports of success in the emotion reappraisal task. Moreover, the interpretation of this null result is complicated by the fact that the fMRI results from the current food choice task differed from previous studies that used similar tasks. In contrast to previous studies (Hare et al., 2009; Hare et al., 2011; Maier et al., 2015; Spetter et al., 2017; van Meer et al., 2017), we did not find significantly increased BOLD activity in the PFC as a function of dietary self-control success.

### Dietary self-control task

Despite not showing any significant increase in PFC as a function of self-control, the participants in the current sample often made the healthier choice when faced with dietary self-control challenges. In fact, the mean overall dietary self-control success level in the current sample is among the highest we have observed across several similar experiments. Even so, the behavioral analysis showed that participants were tempted by highly palatable food items, more often failing to forego eating unhealthy items as they became more tasty. They were also sensitive to the “health cost” of unhealthy foods, being more likely to eat a tasty-unhealthy food if it was relatively less unhealthy (i.e. if the potential negative impact on health was lower). These results indicate that participants remained sensitive to health and tastiness attributes and did not simply follow a rule. Instead, they suggest that participants tried to actively modulate their behavior based on taste and health considerations.

Why, then, does the self-control success BOLD contrast in our dataset differ from the results in previous studies? One potential reason is that, although participants made active goal-directed choices, they also showed a bias toward refusing to eat the food items in terms of both choice outcomes and response times (i.e. faster refuse responses). We presented participants with self-control challenges on approximately 75% of the trials. In challenge trials, participants most often faced decisions in which success required them to refuse palatable-unhealthy foods. A bias toward refusing would facilitate self-control in such cases. Indeed, the mean self-control success level for refusing palatable-unhealthy foods was 77%. Furthermore, within this subset of challenges, successful self-control decisions were actually faster than choices to give into the taste temptations. The correlations between activity in visual processing regions and self-control in our data and previous EEG studies (Harris et al., 2013) also suggest that participants may strategically bias information processing or decision strategies early in, or even prior to, choices in order to facilitate self-control.

Unlike the palatable-unhealthy challenges, self-control success was low in unpalatable-healthy trials. The mean success level for accepting unpalatable-healthy foods was only 19%. Successful healthier choices were slower than failures in this type of challenge as well. This is the opposite of the success-versus-failure response-time pattern seen in palatable-unhealthy challenges.

The difference in self-control success levels between challenge types is consistent with previous reports, but the pattern of response times differs (Hare et al., 2009; Demos et al., 2017). In previous studies, self-control response times were generally slower or not significantly different than decisions that did not present a self-control challenge. However, it is worth noting that the Hare et al. (2009) study upon which the current food choice task was based did not tailor the choice set to each individual and the median percentage of self-control challenge trials was only 22%. In other words, challenge trials occurred relatively rarely in that study, but were common in our current implementation of the task. The frequency of self-control challenges may have led participants to maintain tonic control-related activity in dlPFC and other brain regions. Alternatively, the frequent challenges may have prompted participants to engage phasic regulatory activity at the onset of each trial before determining if regulation was, in fact, needed in the current decision problem. Yet another possibility is that the frequent challenges might have led participants to shift to a decision mode that focused on healthiness attributes without the need for either tonic or phasic control-related brain activity.

Theories of self-control predict that the frequency of self-control challenges will influence the probability of engaging in regulation. The key assumption in these theories is that self-regulation entails some form of costly monitoring and effort that decision makers seek to minimize. Therefore, an individual will use self-control only when the cost of monitoring and trying to influence value computations (i.e. regulating) is smaller than the expected benefit of doing so (Botvinick and Rosen, 2009; Kool et al., 2010; McGuire and Botvinick, 2010; Kool et al., 2013; Shenhav et al., 2013; Shenhav et al., 2017). This calculation depends on how important it is to the decision maker to choose healthy, and the state of the environment. For example, Brocas and Carillo (2019), theorize that in an environment consisting mainly of palatable-unhealthy items, an individual could minimize regulation costs by deciding *a priori* not consume foods unless she detects a healthy option rather than actively regulating on each trial. The pattern of reaction times in our data (faster refusals) is consistent with such a strategy. Notably, all of these theories assume that individuals track what is at stake or the importance of control on each decision.

### Tracking what is at stake in self-control challenges

Therefore, we conducted an exploratory analysis to look for patterns of BOLD activity that correlated with the self-control stake size on each trial. We defined the stakes as the sum of what could be gained and lost in each self-control challenge (see Eq. 5). We found that BOLD activity in prefrontal cortex correlated with the stake size in the current participant sample, and that this result could be replicated when we repeated the same analysis in an independent dataset (Maier et al., 2015). These results suggest that individuals track what is at stake in each self-control challenge as the theories mentioned above predict.

In summary, we cautiously speculate that our task design permitted a simplifying strategy that allowed participants to make healthy choices with less need for choice-specific dlPFC-based regulation or modulation of the value computation process. Specifically, we think that the high frequency of self-control challenge trials together with the high proportion of palatable-unhealthy options within those trials prompted participants to bias their choices toward refusing to eat the proffered food items. This bias to refuse to eat food items may have reduced the need for trial-wise dlPFC engagement. This pattern of behavior may also reflect a shift from reactive to proactive forms of self-control during dietary choice (Braver, 2012; Duckworth et al., 2016). Therefore, we interpret our results from the food choice task as evidence for context-dependent adaptations in self-control strategies. This context-specificity has important implications for the design and utilization of food choice and other paradigms designed to probe self-control and neural activity. However, we emphasize that we can only speculate at this point and further research examining the recruitment of dlPFC for self-control in different choice environments is needed to test these hypotheses more directly.

### Reappraisal task

In contrast to the food choice task, our fMRI results for the emotional stimulus reappraisal task were quite consistent with previous reports on the regulation of responses to affective stimuli (Ochsner et al., 2002; Ochsner and Gross, 2005; Wager et al., 2008; Buhle et al., 2014; Kohn et al., 2014; Morawetz et al., 2017a) or food images (Hollmann et al., 2012; Han et al., 2018). Previous studies of emotion reappraisal have generally focused on the reappraisal of negative scenes and emotional reactions. Here, we extended the emotion reappraisal task to include the regulation of positive affective responses as well. Participants successfully regulated their reactions to both positive and negative stimuli. We did not find any significant differences in BOLD activity during positive versus negative emotion reappraisal. These results suggest that similar systems mediate the reappraisal of both affective valences. However, the standard cautions about (over-)interpreting null results apply to this result as well.

### Limitations and further directions

We wish to point out a few limitations of this study. First, the self-control contrasts for the dietary choice task did not survive correction for multiple comparisons. This precluded computing a conjunction between dietary self-control and emotion regulation success contrasts, which was one of the original objectives of this study. Second, a high frequency of self-control challenges was presented (ca. 75% of all trials). Together with the high proportion of palatable-unhealthy options within those challenge trials, this design feature may have prompted participants to bias the starting point for their choices toward refusing to eat the proffered foods. The ratio of health challenge to non-challenge trials should be carefully considered when designing future studies.

## Conclusion

In conclusion, we found that BOLD activity during emotion reappraisal is positively correlated with dietary self-control. In the case of dietary self-control, we can think of modulating the subjective values placed on the tastiness and healthiness attributes as a modification of the valuation or appraisal process used to place an overall value on the food items. This re- or modified appraisal of the food items leads to healthier choices, which is the goal of dietary self-control in this task. Our findings thus suggest that the neural systems supporting emotion reappraisal can generalize to other behavioral contexts that require reevaluation to conform to the current goal.

## Acknowledgements

The authors thank Astrid Dobler and Jonathan Schaffner for assistance with the data collection and organization and Jan Engelmann, Marcus Grueschow, and Susanne di Pietroantonio for advice on the experiment setup.

## Supplementary Online Material

### Supplementary Methods

#### Emotion regulation success across valence domains

To test whether emotion regulation was equally successful across stimuli with positive and negative valence, we estimated a regression model according to equation S1 below

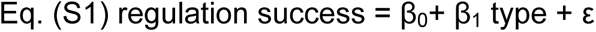

In this model, *regulation success* was defined for negative-valence stimuli as the difference, Reappraisal minus View, because the reappraised rating should be higher (i.e. more positive) than the unregulated viewing rating if reappraisal of negative stimuli was successful. The difference, View minus Reappraise, described success in the positive reappraisal trials, because for positive stimuli the unregulated View ratings should be higher than the reappraised rating when successfully modulating positive emotions. Trial *type* was a factor with 2 levels (1 = negative, 2 = positive valence). The model included subject-specific random intercepts and slopes for the trial type.

#### Drift diffusion modeling

In order to test whether there was a decision bias towards refusing foods, we modeled the data using a drift diffusion model that allows for different attribute onset times for taste and health. For full details on this modeling approach, we refer the reader to the paper by Maier et al. (in press, preprint available at bioRxiv (Maier et al., 2020)). Briefly, this model is a time-varying drift diffusion model (DDM) that is augmented by one parameter that captures the start time of health attribute processing relative to the processing start time of taste (parameter *RST*). This feature of the model allows for better identification of the weight quantifying the influence of each attribute on the evidence accumulation process and any starting-point bias. In our formulation of the model, a negative sign on the bias signifies that participants had a starting point bias in favor of refusing to eat the food.

We estimated this model on the food choices in the current dataset (N=39 participants who are included in the dietary self-control analyses, hereafter abbreviated as study *ESC*) and used two-sample BEST tests (Kruschke, 2013) to compare the starting-point bias (hereafter in short: *Bias*) and relative-health-start-time (*RST*) estimates to the results of (Hare et al., 2011) (hereafter abbreviated as study *IAC*). This study was employing the same choice setup of choosing the food on the screen versus nothing and also had a condition with a health reminder.

#### Reaction time simulations

In order to test whether reaction times generated by the DDM could yield the pattern of results observed in Table 4, we simulated reaction times for all trials using the best generating DDM parameters and the taste and health value difference from zero (i.e., when refusing to eat) for each participant.

Corresponding to the choice boundary definition that was used in the fitting of the DDM, we recorded the corresponding simulated choices as “yes” if they had a positive sign, and as “no” if they had a negative sign. We took the absolute value of the simulated RTs and applied the natural logarithm in order to fit them using the regression model described in Eq. 4.

#### Correlations between individuals’ BOLD responses and their DDM Bias parameters

In order to test for correlations between BOLD responses and the Bias parameter, we ran a regression using all 6 DDM parameters and an intercept, because the DDM parameters are interdependent. Within this group-level regression model, we specified two contrasts testing for either a positive or a negative correlation with the Bias parameter. We tested whether the individual levels of these parameters explained individual variance in the BOLD signal of 1) the contrast “Self-Control Success > Self-Control Failure” in GLM-SCS and 2) of the contrast “All Choice” in GLM-FC.

### Supplementary Results

#### Drift diffusion model parameter comparison to previous results

We first tested whether the participants of the current study (ESC) expressed a bias towards refusing the foods. The results in Table S2 show a negative sign for the bias term, indicating that there was indeed a bias towards refusing the foods in the current study. The comparison with the study of Hare et al. (2011) showed that compared to the health-cue condition in IAC, participants in the current study expressed a greater bias towards saying “No” (difference in starting point bias = −0.156, Posterior Probability (PP) of (ESC bias < IAC health condition bias) = 0.98, 95% HDI = [-0.30; −0.01]). Note that, in our model formulation, negative bias parameters favor the “No” response. As expected, the difference in the bias terms across studies was even more pronounced when comparing to the natural choices in IAC (difference in starting point bias = −0.377, PP(ESC bias < IAC natural condition bias) = 0.999, 95% HDI = [-0.54; −0.22]). This indicates that across all trials in the present study participants showed a greater inclination towards refusing the foods, and that this bias was greater than could have been expected from the comparable study.

We also tested whether there were any differences in the attribute consideration onset timing (RST). Here, we did not observe differences between the ESC sample and the IAC health condition, which is most comparable to our present study setup where we also introduced a health reminder. The relative start time for health did not differ significantly between the ESC study and the IAC health condition: the mean difference in the RST parameter was −0.04 (PP(RST ESC > RST IAC health condition) = 0.39, 95% HDI = [-0.28; 0.21]).

#### Simulation results from the DDM

In order to test whether an overall bias toward responding “Not Eat” – which is present in the DDM parameters – can still lead to the reaction time pattern we observe in Figure 3, we simulated reaction times based on the best-fitting DDM parameters for each subject. The simulated RTs yielded a very similar pattern. Reaction times were faster when the simulated agents refused to eat the foods, except for the palatable-healthy foods where accept response was favored by the healthiness and palatability attributes (Supplementary Figure S2). These simulation results indicate that our empirical results are consistent with an overall bias toward refusing to eat the foods.

#### Neural correlations with DDM Bias

Testing for neural correlates of the DDM Bias parameters yielded no results that survived whole-brain correction (Table S4).

#### Behavioral associations between tasks

At an anonymous reviewer’s request, we tested whether there were behavioral associations between the tasks when correlating the task performance within the domain of appetitive / positive and aversive / negative stimuli.

The analysis in separate domains did not yield significant correlations either. For the correlation of positive emotion regulation success with the overall success level in refusing to eat palatable-unhealthy foods, Spearman’s rho was 0.19, Posterior Probability(Rho > 0) = 0.86, 95% HDI = [-0.13; 0.51]. For the correlation of the negative emotion regulation success with the overall success level in accepting to eat healthy-unpalatable foods, Spearman’s rho was −0.16, PP (Rho < 0) = 0.82, 95% HDI = [-0.50; 0.18].

## Supplementary Figures

**Figure S1.**
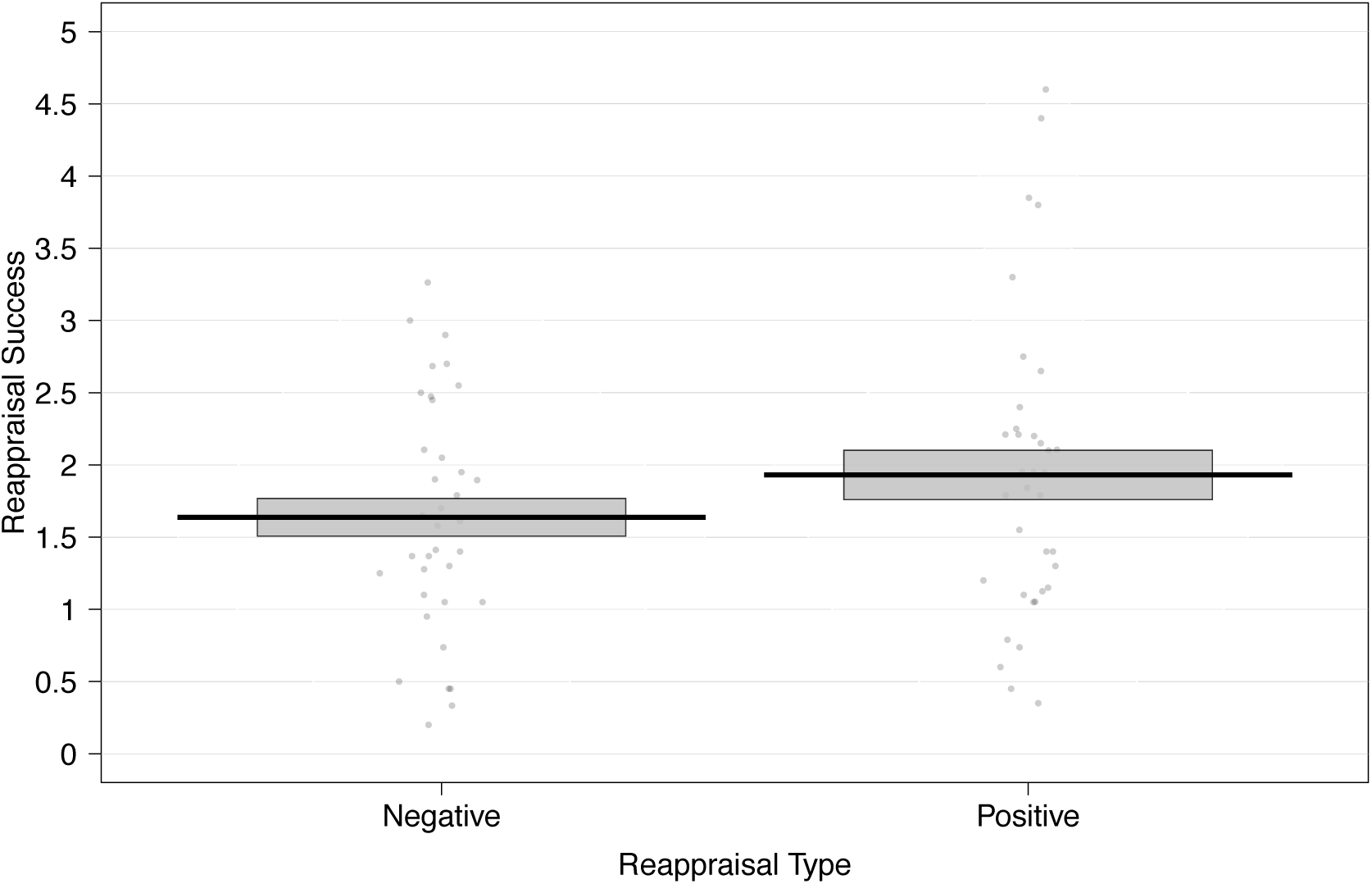
Reappraisal Success by trial type. This figure shows the mean reappraisal success for both types of reappraisal. Reappraisal success was the trial-wise difference of the rating given after regulation minus the rating given when just viewing the stimulus for negative stimuli, and vice versa for positive stimuli. The black solid line represents the group mean and the gray box indicates the standard error of the group mean. Each dot represents the mean reappraisal success by reappraisal type for one participant.

**Figure S2.**
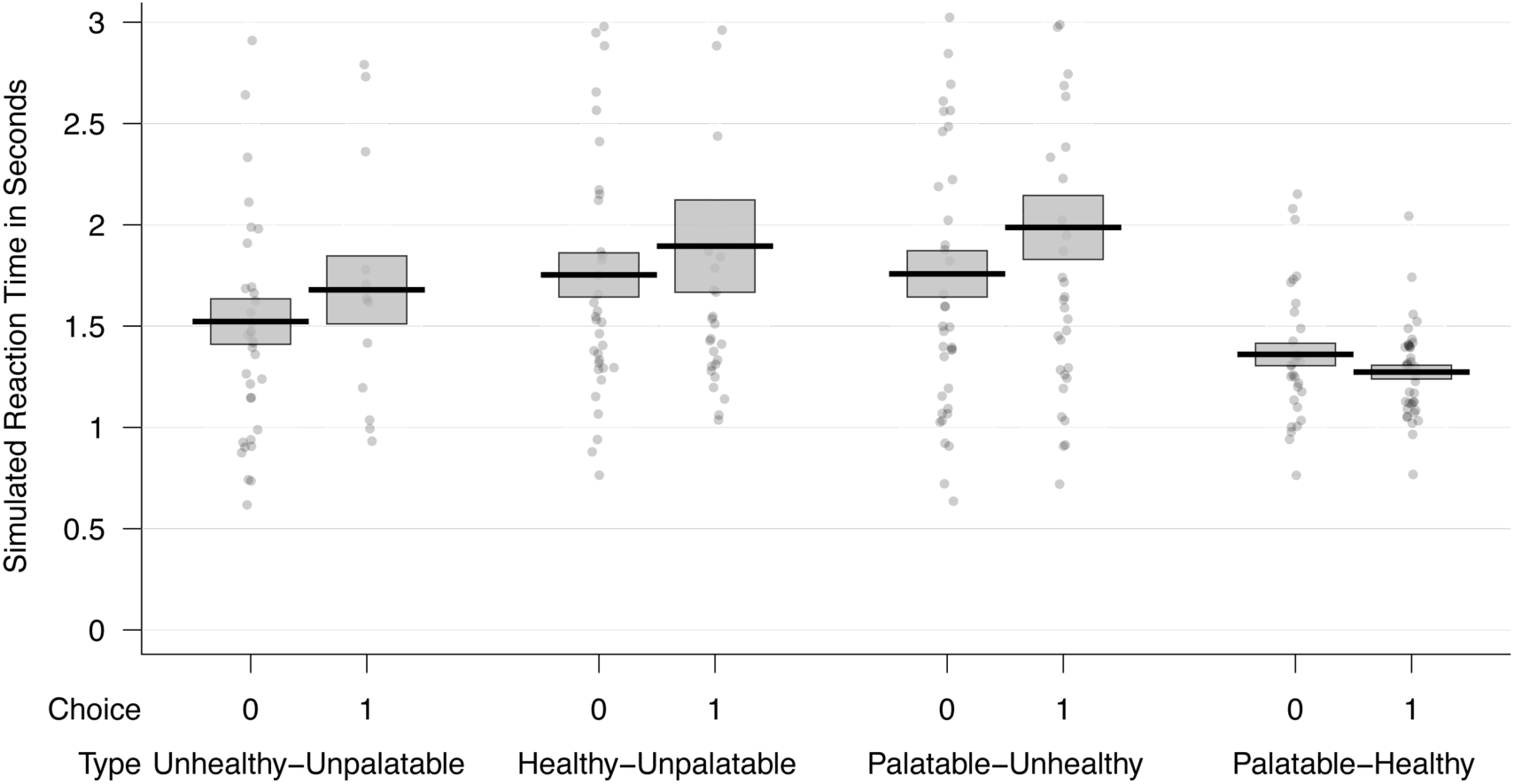
Simulated reaction times generated from the best-fitting DDM parameters (analogous to Figure 3 in the main text). This figure shows the mean reaction times (RTs) over all simulated participants for accepting (Choice = 1) and refusing (Choice = 0) to eat foods from each of the four categories. The black solid line represents the group mean and the gray box indicates the standard error of the group mean. Each dot represents the mean RT by choice category for one simulated participant. On average, the generating drift diffusion models, on which each participant’s simulations were based, had a starting point bias towards refusing to eat the foods. Qualitatively, this plot of the simulated reaction times for the different food categories captures the features of Figure 3 in the main text. This suggests that the results observed in the experiment are consistent with a starting-point-bias in favor of refusing the foods.

**Figure S3.**
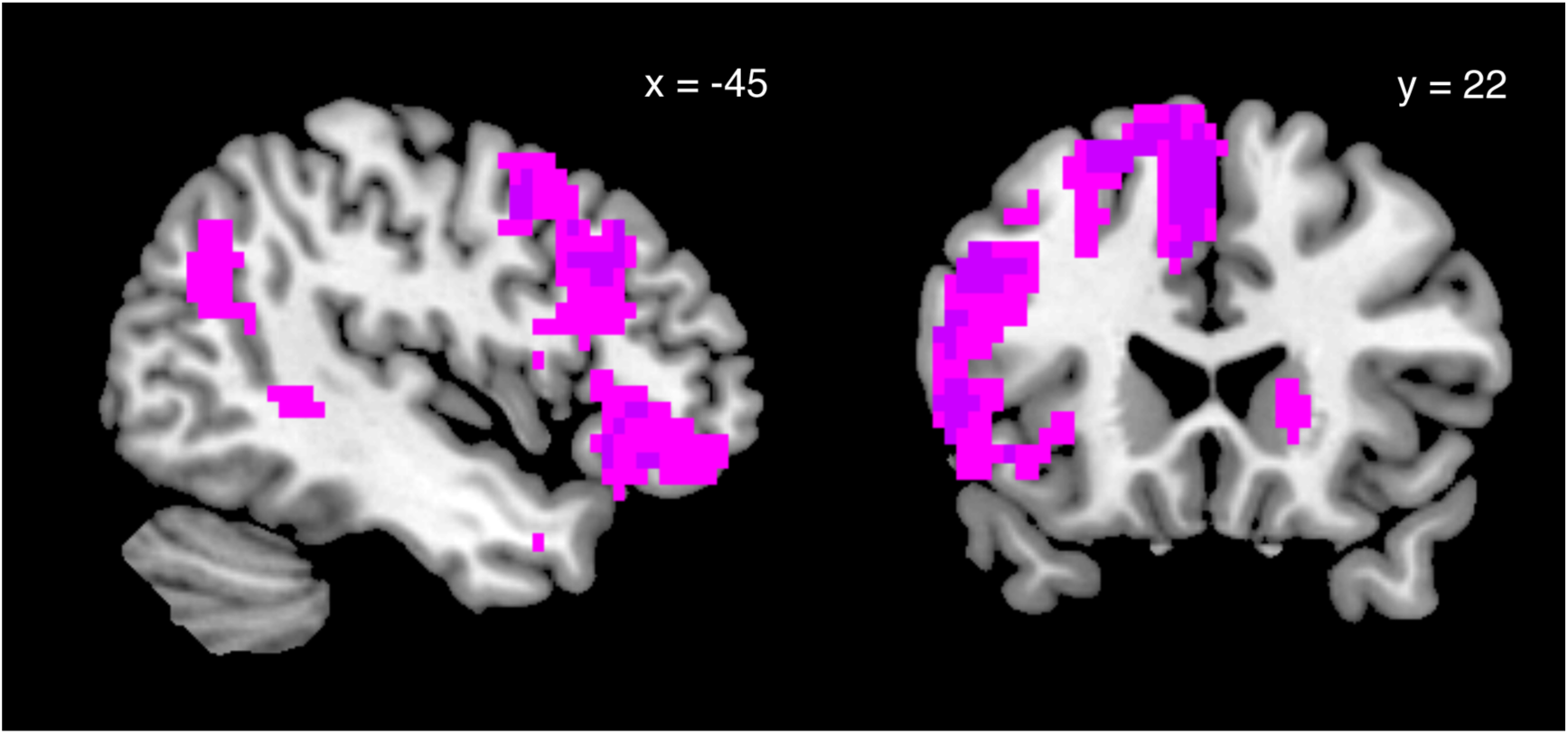
Whole-brain family-wise error corrected group-level results (p < 0.05) for the contrast Reappraise > View (violet, darker shading) and the contrast Reappraisal Success > View (pink, lighter shading) from GLM-ER. The two contrasts are very similar. This is not surprising because participants rarely failed to reappraise the image content.

## Supplementary Tables

**Table S1.**
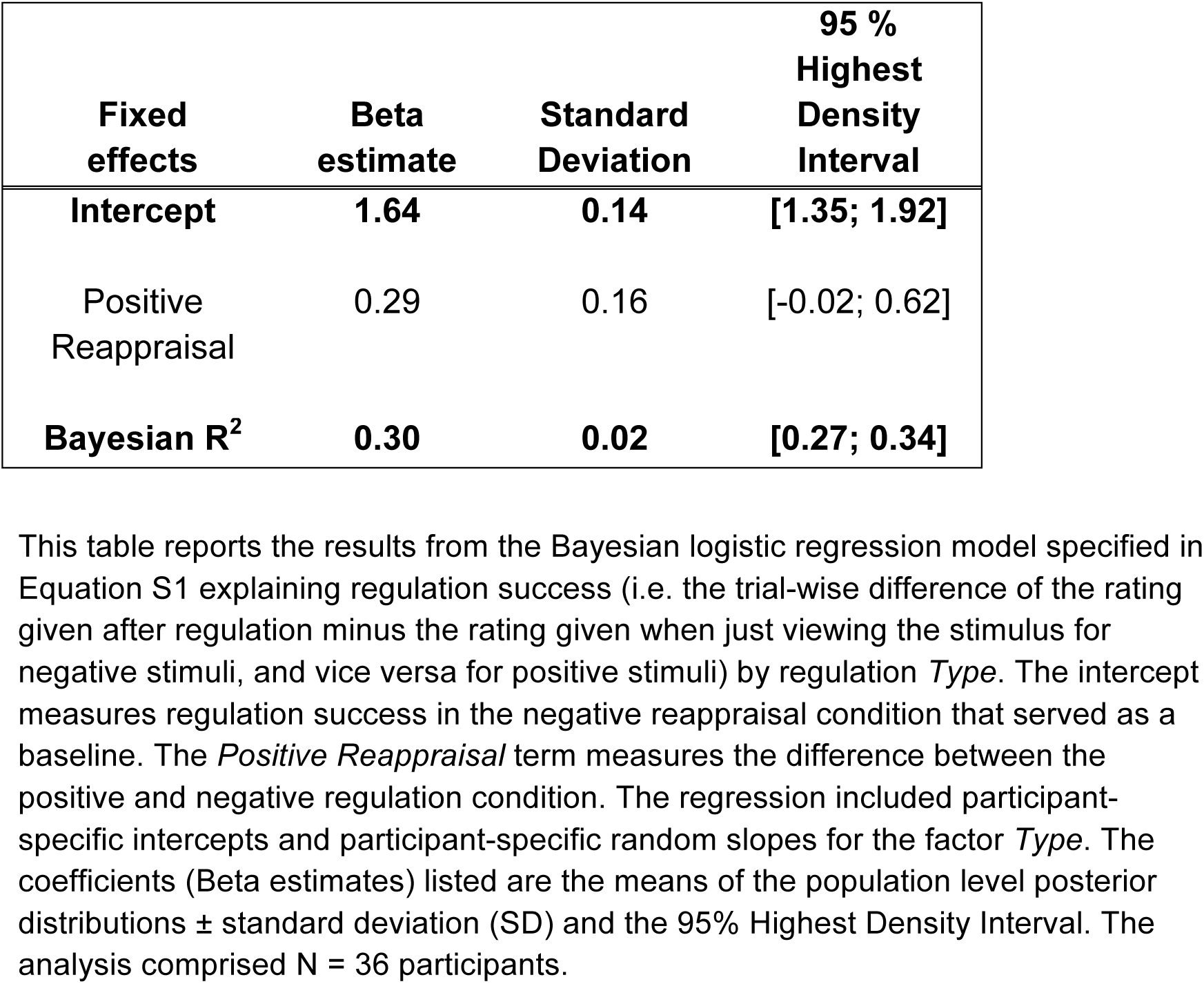
Regression modeling positive versus negative reappraisal success.

| Fixed effects | Beta estimate | Standard Deviation | 95 % Highest Density Interval |
| --- | --- | --- | --- |
| Intercept | 1.64 | 0.14 | [1.35; 1.92] |
| Positive Reappraisal | 0.29 | 0.16 | [-0.02; 0.62] |
| Bayesian $R^2$ | 0.30 | 0.02 | [0.27; 0.34] |

This table reports the results from the Bayesian logistic regression model specified in Equation S1 explaining regulation success (i.e. the trial-wise difference of the rating given after regulation minus the rating given when just viewing the stimulus for negative stimuli, and vice versa for positive stimuli) by regulation *Type*. The intercept measures regulation success in the negative reappraisal condition that served as a baseline. The *Positive Reappraisal* term measures the difference between the positive and negative regulation condition. The regression included participant-specific intercepts and participant-specific random slopes for the factor *Type*. The coefficients (Beta estimates) listed are the means of the population level posterior distributions ± standard deviation (SD) and the 95% Highest Density Interval. The analysis comprised N = 36 participants.

**Table S2.** . Time-varying DDM parameters.

| Dataset | Parameter estimate |  |  |  |  |  |
| --- | --- | --- | --- | --- | --- | --- |
| (1) *ESC* | $\omega_{\text{taste}}$ | $\omega_{\text{health}}$ | Thr | nDT | RST | Bias |
| Current food choices | 0.36 ± 0.40 | 0.41 ± 0.26 | 1.13 ± 0.27 | 0.72 ± 0.12 | -0.02 ± 0.43 | -0.37 ± 0.29 |
| (2) *IAC* | $\omega_{\text{taste}}$ | $\omega_{\text{health}}$ | Thr | nDT | RST | Bias |
| Natural Choice | 1.37 ± 0.79 | 0.33 ± 1.33 | 1.27 ± 0.28 | 0.86 ± 0.12 | 0.42 ± 0.54 | 0 ± 0.37 |
| Health Cued choices | 0.98 ± 1.12 | 1.11 ± 0.60 | 1.39 ± 0.36 | 0.85 ± 0.14 | -0.06 ± 0.55 | -0.22 ± 0.33 |

This table reports the group mean ( ± standard deviation) of the best-fitting parameters from a time-varying DDM model with separate attribute consideration onset times. The rows report the data from the current dietary self-control task (current food choices) and both the Health-cued and Natural (i.e. baseline) choices from a similar study by Hare et al. (2011) in which participants were given instructed attention cues (IAC) toward healthiness or palatability. Abbreviations:
**ESC:** the food choice data from the current study (Emotion regulation and dietary self-control). All food choices in this study were made under the instruction to consider healthiness and try to make healthy choices.
**IAC:** the food choice data from the instructed attention cue study by Hare et al. (2011). In the Health Cue" condition, participants were asked to focus on the health attributes of the presented food when making their choices, whereas in the "Natural Choice" condition, they were asked to choose as they naturally would do.
$\omega_{\text{taste}}$ = weighting factor determining how much the difference in taste attributes contributes to the evidence accumulation rate.
$\omega_{\text{health}}$ = weighting factor determining how much the difference in health attributes contributes to the evidence accumulation rate.
Thr = evidence threshold for responding.
nDT = non-decision time and corresponds to the starting time for taste in our model.
RST: relative start time for health (timing relative to start of taste processing, positive values denote that health enters the process later than taste).
Bias: starting point bias for the evidence accumulation process (zero = no bias).

**Table S3.** Results of the Reappraisal > View contrast from GLM-ER.

| *Region* | *Side* | *MNI Coordinates* |  |  | *TFCE t-stat* |
| --- | --- | --- | --- | --- | --- |
| Superior Frontal Gyrus | L | -5 | 12 | 61 | 6.07 |
| Inferior Frontal Gyrus, pars triangularis | L | -53 | 27 | 18 | 5.02 |
| Middle Frontal Gyrus | L | -35 | 4 | 61 | 5.05 |
| Middle Frontal Gyrus | L | -48 | 24 | 32 | 5.81 |
| Middle Temporal Gyrus, posterior division | L | -53 | -44 | 0 | 6.8 |
| Superior Frontal Gyrus | L | -5 | 47 | 43 | 6.24 |
| Middle Frontal Gyrus | L | -45 | 17 | 40 | 4.29 |

**Table S4.** . BOLD signals were not significantly correlated with starting point biases.

| **(1) Self-Control Success > Self-Control Failure** | **DDM parameter** | **Direction of correlation** | **Minimum p-value** |
| --- | --- | --- | --- |
|  | Bias | positive | 0.25 |
|  | Bias | negative | 0.83 |
| **(2) All Food Choices** |  |  |  |
|  | Bias | positive | 0.50 |
|  | Bias | negative | 0.85 |

We conducted a set of exploratory analyses to see if starting-point bias parameters from the DDM correlated with BOLD activity in any voxels at the time of choice. We used two contrasts, 1) Self-Control Success > Self-Control Failure contrast from GLM-SCS, and 2) the All Choice contrast from GLM-FC. There were no significant correlations after correcting for multiple comparisons at the whole brain level. Here, we report the brain-wide minimum p-value for each contrast and correlation direction. The correlations with *All Food Choices* comprise N = 37 participants. The correlations with *Self-Control Success > Self-Control Failure* comprise N = 35 participants because this contrast could not be evaluated for two participants due to too few failures.

